# Proteasome inhibitor-induced modulation reveals the spliceosome as a specific therapeutic vulnerability in multiple myeloma

**DOI:** 10.1101/508549

**Authors:** Hector H. Huang, Ian D. Ferguson, Alexis M. Thornton, Christine Lam, Yu-Hsiu T. Lin, Prabhakar Bastola, Priya Choudhry, Margarette C. Mariano, Makeba Marcoulis, Julia Malato, Paul Phojanakong, Thomas G. Martin, Jeffrey L. Wolf, Sandy W. Wong, Nina Shah, Byron C. Hann, Angela N. Brooks, Arun P. Wiita

## Abstract

Enhancing the efficacy of proteasome inhibitors is a central goal in myeloma therapy. We proposed that signaling-level responses after PI would reveal new mechanisms of action that could be therapeutically exploited. Unbiased phosphoproteomics after the PI carfilzomib surprisingly demonstrated the most prominent phosphorylation changes on splicing related proteins. Spliceosome modulation was invisible to RNA or protein abundance alone. Transcriptome analysis after PI demonstrated broad-scale intron retention, suggestive of spliceosome interference, as well as specific alternative splicing of protein homeostasis machinery components. These findings led us to evaluate direct spliceosome inhibition in myeloma, which synergized with carfilzomib and showed potent anti-tumor activity. Functional genomics and exome sequencing further supported the spliceosome as a specific vulnerability in myeloma. Our results propose splicing interference as an unrecognized modality of PI mechanism, reveal additional modes of spliceosome modulation, and suggest spliceosome targeting as a promising therapeutic strategy in myeloma.

**Significance:** New ways to enhance PI efficacy are of major interest. We combine systems-level analyses to discover that PIs specifically interfere with splicing and that myeloma is selectively vulnerable to spliceosome inhibition. We reveal a new approach to advance myeloma therapy and uncover broader roles of splicing modulation in cancer.

## INTRODUCTION

Multiple myeloma is a clonal malignancy of plasma cells with no known cure. Like normal plasma cells, myeloma cells produce and secrete incredible amounts of immunoglobulin. This unique function may be exploited by therapeutically inhibiting the proteasome using the FDA-approved proteasome inhibitors (PIs) bortezomib, carfilzomib, and ixazomib. Proteotoxic stress caused by these first-line therapeutic agents has been proposed to induce the apoptotic function of the unfolded protein response (UPR) (1), leading to plasma cell death while largely sparing normal tissues (2, 3). However, despite the appealing simplicity of this mechanism, the canonical UPR is not always strongly induced in myeloma cells by PIs (4) and is unlikely to be the sole mode of PI cytotoxicity in MM. Indeed, many additional mechanisms of action of PIs have also been proposed, ranging from NF-kB inhibition to immune microenvironment effects to aberrant recycling of cytosolic amino acids (5, 6).

Identifying the full range of PI mechanisms of action remains relevant given that acquired PI resistance is clinically widespread but its origins remain unclear (7, 8) and finding new methods to specifically target PI-resistant disease, or molecules to synergize with PIs to avoid resistance by driving deeper remissions, remains a long-standing goal. As one approach to achieving this goal, we and others have studied the response of malignant plasma cells to PIs using both gene expression and proteomic methods (9-11). Notably, one of the most prominent features of the cellular response to PIs is the activation of the heat shock response (12). This mechanism leads to significant induction of cytosolic protein-folding chaperones, possibly to assist in protein refolding and decrease in unfolded protein stress. We and others (9, 12, 13) have therefore proposed targeting mediators of the heat shock response as potential combination therapies with PIs.

However, one unresolved question is whether proteasome inhibition may carry additional effects on plasma cells that are not revealed by mRNA or protein abundance analysis alone. We hypothesized that additional modalities of response, and thereby new myeloma-relevant therapeutic targets, may be revealed by studying the signaling-level response to PIs with unbiased mass spectrometry-based phosphoproteomics. The large majority of therapy-relevant investigations using this technique have focused on elucidating the effects of kinase inhibitors (14). However, we reasoned that a significant cellular perturbation such as proteasome inhibition would likely also indirectly perturb kinase and phosphatase signaling in a broad fashion.

Here, we used unbiased phosphoproteomics to quantify >5000 phosphopeptides in myeloma cells exposed to the irreversible PI, carfilzomib (Cfz). Surprisingly, we found the greatest increases in phosphorylation occurred in proteins associated with the spliceosome machinery. A link between these processes was invisible at the gene expression level. We further evaluated this link from a mechanistic and therapeutic perspective, finding that PIs lead to specific disruption of normal splicing. We suggest interference of splicing as an additional mechanism of action of PIs not previously explored. Inhibition of splicing has recently become a promising therapeutic strategy in other hematologic malignancies (15). Our results reveal an intersection of cellular stress and the splicing machinery, which may have broad relevance in biology. Furthermore, we propose the spliceosome as a new and potentially selective therapeutic target in myeloma.

## RESULTS

### Proteasome inhibition results in sustained phosphorylation of splicing factors in myeloma plasma cells

We first used unbiased phosphoproteomics to examine the signaling-level response of MM.1S multiple myeloma cells to Cfz. We chose time points across 24 hours for analysis based on our prior results demonstrating that the transcriptional and proteomic response to proteasome inhibition evolves over many hours (9). This is in contrast with the majority of prior perturbation phosphoproteomic studies, which have typically examined direct effects on kinase activation or inhibition on a timescale of minutes (14). Here, we instead consider the indirect effects on phosphorylation induced by PI exposure. Using label-free quantification of immobilized metal affinity chromatography (IMAC)- isolated phosphopeptides, we indeed found that altered phosphorylation signatures were most prominent 24 hours after treatment (**Fig. 1 and Fig. S1**). In total, we quantified 5791 phosphosites in at least one technical replicate of the time course, with >99% of phosphosites representing Ser or Thr phosphorylation events, as expected using this enrichment technique. Notably, with 30 nM Cfz, cell viability was approximately 30% of baseline, indicating significant drug-induced cytotoxicity by this final time point.

**Figure 1.**
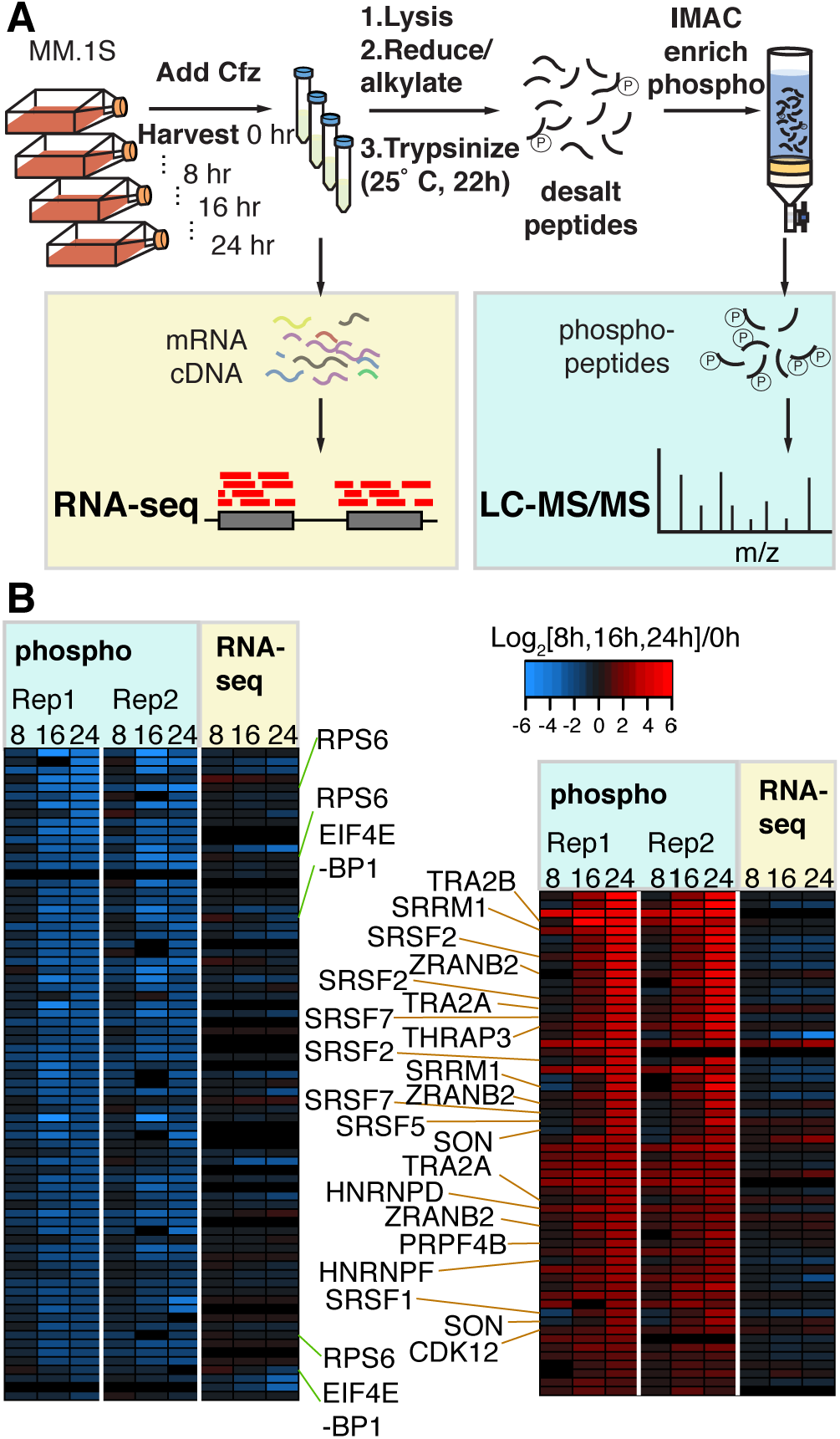
Unbiased phosphoproteomic timecourse analysis of MM.1S cells treated with the PI carfilzomib (Cfz). **A.** Workflow of timecourse treatment of MM.1S cells with Cfz. Cells were allotted for both RNA-seq analysis and LC-MS/MS. **B.** Downregulated (left) and upregulated (right) log_2_ transformed phospho-peptide MS1 intensities for two technical replicates of proteins with unchanged transcript levels (RNA-seq). Labels highlight dephosphorylation of RPS6 and EIF4EBP1 on the left and phosphorylation of splicing-related proteins on the right.

At each time point we simultaneously performed single-end RNA-seq to determine gene expression to compare with our phosphoproteomic results. **Fig. 1B** shows 58 upregulated (red) and 75 downregulated (blue) phosphopeptides from proteins with largely unchanged RNA transcript abundance as detected by unsupervised hierarchical clustering. Upon this initial analysis, we were encouraged to find decreased phosphorylation of the translation factor EIF4E- BP1 as well as the ribosomal subunit RPS6 (**Fig. 1B**). These phosphorylation-level responses related to suppressed translation are expected upon PI-induced cellular stress (9). While other downregulated phosphopeptides did not suggest a specific highly-enriched biological function, upon manual inspection of upregulated phosphosites we were surprised to find that 14 of 58 were present on proteins related to pre-mRNA splicing. These primarily included phosphopeptides deriving from the heterogeneous ribonucleoprotein (HNRNP) family of proteins as well as phosphopeptides belonging to the SRSF family of splicing factors (**Fig. 1B**). In particular, the arginine- and serine-rich “RS” domain of the SRSF proteins are known to have their splicing activity modulated by phosphorylation (16). Notably, these prominent signaling-level effects on splicing factors were invisible to prior gene expression studies of PI response and have not been investigated previously. We therefore chose to further explore the interaction between PIs and the splicing machinery.

To validate this initial result from label free quantitative proteomics, we prepared independent samples using a stable isotope labeling (SILAC) phosphoproteomics approach. Based on our results above, we examined only the 24 hr time point in MM.1S cells. We evaluated both a low dose (10 nM, *n* = 2 biological replicates) and a moderate dose (18 nM, *n* = 2) of Cfz (**Fig. 2A-B**) With this lot of Cfz, 10 nM drug elicited ∼20% cell death after 24 hr, while 18 nM killed ∼85% of cells (**Fig. S3A**). Of the 520 phosphosites significantly (*p* < 0.05; ≥ 2-fold-change) upregulated in MM.1S treated with 18 nM Cfz in **Fig. 2A**, 127 (24.4%) are associated with splicing-related proteins, with 23 of these as part of the SRSF protein family of splicing factors. Background-corrected Gene Ontology (GO) analysis confirms that all of the top enriched biological processes involve RNA splicing regulation and mRNA processing (**Fig. S1B, 2E**). At 10 nM Cfz, though, this signaling response is much weaker with only 25 upregulated phosphosites; none of these are splicing-related. These results suggest that there is a strong dose-response effect of phosphorylation changes after proteasome inhibition, both across splicing factors and the broader proteome.

**Figure 2.**
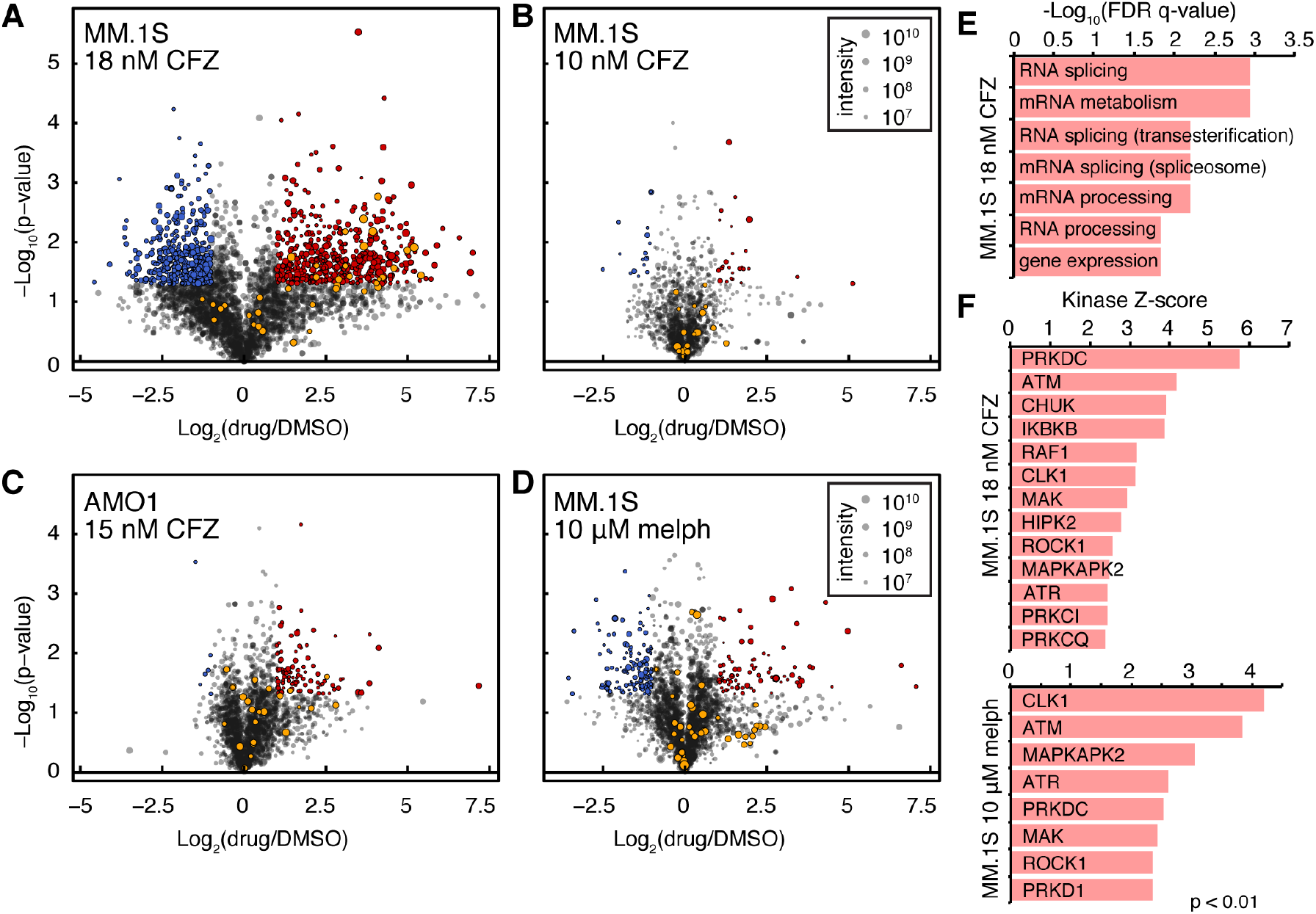
Cfz induces phosphorylation of splicing factors in a dose-responsive manner. **A-D.** Volcano plots of log2 transformed ratios of phosphosite abundances between **A.** MM.1S treated with18 nM Cfz, **B.** 10 nM Cfz, or **D.** or 10 µM melphalan compared to DMSO and **C.** AMO-1 with 15 nM Cfz compared to DMSO. Significant upregulated sites are in red, while downregulated are in blue (>2-fold change, *p* < 0.05). SRSF related sites are in orange. Circle size corresponds to summed SILAC intensities. **E.** Top-ranked GO terms for genes with significantly upregulated phosphosites in MM.1S cells treated with 18 nM Cfz. **F**. Top-ranked KSEA activated kinases (with at least 5 substrates) for MM.1S treated with 18 nM Cfz (top) and 10 µM melphalan (bottom).

To compare these changes at the signaling level to changes at the protein level, unenriched peptides were also analyzed by LC-MS/MS (**Fig. S2A-B**). Confirming expected responses to proteasome inhibition, the most upregulated proteins included heat shock-induced chaperones as well as SQSTM1/p62 associated with autophagy (9). In contrast, splicing factors with increased phosphorylation sites do not significantly change in abundance, confirming that phosphosite increases are due to changes at the signaling level and not protein copy number.

### Melphalan induces a similar but not identical phosphorylation response

We next investigated whether this broad splicing factor phosphorylation phenotype was unique to proteasome inhibition or was also seen under a different drug mechanism of action. We chose to compare the response of MM.1S cells to melphalan, a DNA alkylating agent and clinically-used myeloma therapeutic. In parallel, we also treated another MM cell line, AMO-1, with Cfz to determine if the phosphorylation response to proteasome inhibitor is consistent across cell line models.

For these experiments we again used a single-timepoint SILAC approach. Here, both 10 µM melphalan and 15 nM Cfz led to ∼20% cell death in MM.1S and AMO-1, respectively, at 24 hr (**Fig. S3A**). Western blot confirmed induction of DNA damage by melphalan and proteotoxic stress response for Cfz (**Fig. S3C-D**). Compared to 18 nM Cfz, we saw largely decreased phosphorylation-level responses to both of these agents (**Fig. 2C-D**). Of 113 phosphosites significantly upregulated in AMO-1, 7 belong to splicing related proteins (SRSF2, SRSF6, SRRM1, HNRNPH1, TRA2A, DDX1). This result is consistent with the MM.1S results in **Fig. 2A-B**, where greater PI response correlates with more prominent phosphorylation changes.

Under 10 µM melphalan, 93 phosphosites were significantly upregulated, with 8 sites on splicing related proteins (HNRNPK, TRA2A, SRRM2, and WDR77), although none are SRSF family members (**Fig. 2D**). Furthermore, as expected, both unenriched shotgun proteomics and RNA-seq for gene expression confirm that proteasome inhibition and DNA damage elicit different responses (**Fig. S2A-E**). Again, no splicing factors with altered phosphosites under either drug treatment were changed at the protein abundance level.

We further performed Kinase Set Enrichment Analysis (KSEA) (17) on our MM.1S datasets to identify kinases whose activity may regulate differential phosphorylation found by phosphoproteomics. While this tool is limited by its reliance on well-characterized kinase-substrate relationships, and despite the different number of phosphosites upregulated under each condition, within this framework this tool identified similar kinases active under both 18 nM Cfz and 10 µM melphalan treatment (**Fig. 2F**). Notably, both drugs are predicted to induce activity of cdc2-like kinase 1 (CLK1), a kinase known to phosphorylate SRSF family splicing factors among other proteins (18). However, in line with the specific biology of PIs, Cfz also strongly induced inhibitory kappa B kinase (IKBKB) activity, a kinase leading to NF-kB inhibition after PI treatment (19). Taken together, these results indicate that drug-induced stress may broadly lead to phosphorylation of splicing factors, though precise patterns of phosphorylation may differ in a drug-specific manner.

### SRSF splicing factors appear highly phosphorylated at baseline in MM cells

To investigate our phosphoproteomic results via an orthogonal method, we performed Western blots to evaluate for phosphorylation-induced gel mobility shift after Cfz treatment of SRSF1, SRSF3, and SRSF6. After Cfz treatment and isolation of the cytoplasm, we initially saw no discernable shift of these proteins. However, treatment of lysate with calf alkaline phosphatase resulted in a substantial shift of SRSF proteins but not actin (**Fig. S3F**). Therefore, these SRSF factors exist in a highly phosphorylated state even at baseline in MM plasma cells. Upregulated phosphorylation post-Cfz identified by mass spectrometry may therefore represent additional phosphorylation at only selected phosphosites. While these changes in phosphorylation may still result in biological effects, Cfz-induced modulation does not appear to reflect a dramatic shift in the overall phosphorylation status of these SRSF proteins in this system.

To further investigate baseline phosphorylation status of SRSF proteins, we treated MM.1S cells with 50 µM KH-CB19 (20), a reported highly selective inhibitor of the SRSF kinases CLK1 and CLK4 (K_D_ = 20 nM vs. CLK1). We did not observe any viability effects in MM.1S even at this high concentration (**Fig. S3A**). Unbiased phosphoproteomics after 24 hr of KH-CB19 treatment surprisingly showed no significant change in phosphorylation status of any quantified SRSF phosphosites, except one upregulated (**Fig. S2F**). These results suggest that other kinases also play a role in maintaining SRSF phosphorylation in this system, either at baseline or via feedback mechanisms after sustained CLK1 inhibition.

### Proteasome inhibition induces intron retention in MM cells

Given our results demonstrating splicing factor phosphorylation, we next investigated whether pre-mRNA splicing itself was altered after drug treatment. We obtained paired-end sequencing data from polyA-enriched RNA on the same samples used for phosophoproteomics, plus one additional biological replicate (*n* = 3 total) of each of the following: MM.1S treated with 18 nM Cfz, with 10 µM melphalan, and with DMSO as control; and AMO-1 treated with 15 nM Cfz and with DMSO as control. We used JuncBASE (21) to process the aligned sequencing data by identifying and quantifying both annotated and novel splice junctions. Data for each alternative splicing event was evaluated using the standard measure of “percent spliced in” or PSI (ψ) (**Fig. 3A**).

**Figure 3.**
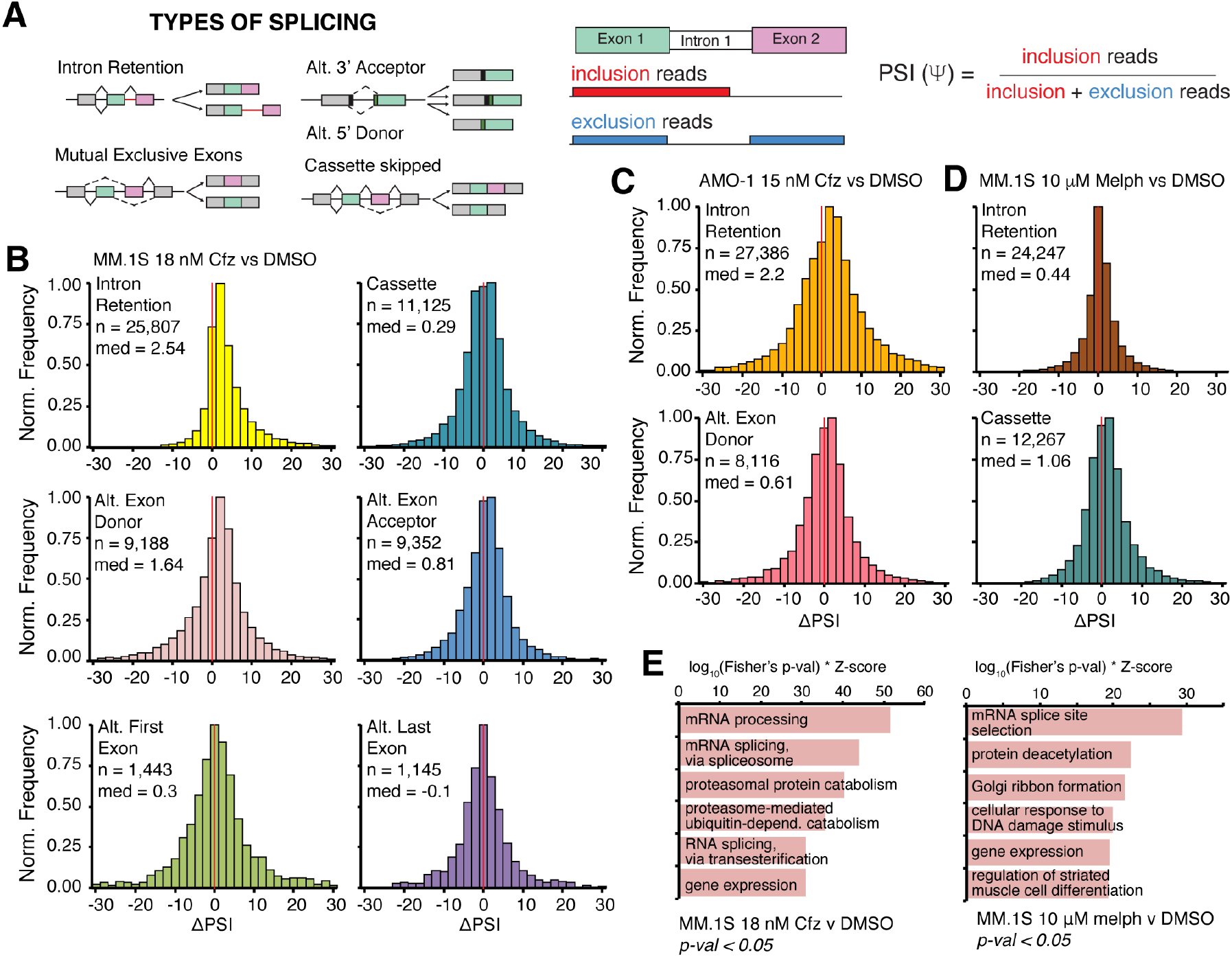
Cfz treatment leads to prominent intron retention. **A.** Cartoon description of alternative splicing event (ASE) types and description of ΔPSI. **B.** Histograms of ΔPSI for JuncBASE identified ASEs in MM.1S treated with 18 nM Cfz stratified according to type of splicing event (IR = yellow, alt. exon cassette = teal, alt. exon donor (5’ splice site) = pink, alt. exon acceptor (3’ splice site) = blue, alt. first exon = green, alt. last exon = purple). Bin = 2, red line indicates ΔPSI = 0. **C.** Histograms of ΔPSI for all IR events (top) and Alt. Exon Donor events (bottom) in AMO-1 treated with 15 nM Cfz **D.** Histograms of ΔPSI for all IR events (top) and Alt. cassette events (bottom) in MM.1S treated with 10 µM melphalan (right). **E.** Top ranked GO enrichment terms for genes with significant (*p* < 0.05) ASEs for MM.1S cells treated with 18 nM Cfz (left) or with 10 µM melphalan (right).

Comparative analysis of differential PSI (ΔPSI) between 18 nM Cfz- and DMSO-treated MM.1S were considered according to categories including alternative exon acceptor (3’ splice site selection), alt. donor (5’ splice site selection), alt. last exon, alt. first exon, alt. exon cassette, and intron retention (IR) (**Fig. 3B, Supplementary Table S3**). The ΔPSI distribution for IR demonstrated the greatest positive shift after Cfz treatment (*n* = 25,807 total IR events measured; median = 2.54). ΔPSI medians for alternative splice site selection also demonstrated a significant shift (alt. donor median = 1.64, *n* = 9,188 and alt. acceptor median = 0.81, *n* = 9,352). All other categories were closer to a median of zero (*p* < 2.2E-16 for median of IR distribution, alt. donor, and alt. acceptor vs. median of cassette by Mann-Whitney test, **Supplementary Table S3**). Intriguingly, PIs are well known to induce a strong heat shock response (12) and prior work in non-cancer cells demonstrated that heat shock alone could impair splicing and induce IR without broadly affecting other alternative splicing events (22, 23). In general, intron-retained transcripts may be subject to nonsense-mediated decay or retained in the nucleus where they remain untranslated. Our results suggest that a similar splicing impairment may be present in MM cells exposed to PI.

We then considered the possibility that the IR phenotype results from a global dysfunction of the splicing machinery during drug-induced apoptosis, which is likely occurring with ∼85% cell death at our high-dose Cfz treatment. Our prior proteomic data indicated that SF3B1 and U2AF2, core components of the splicing machinery, are some of the earliest substrates cleaved by caspases during PI-induced apoptosis (9). Indeed, by Western blotting we validated that SF3B1 and U2AF2 are proteolytically cleaved after Cfz treatment and this cleavage can be blocked by the pan-caspase inhibitor zVAD-fmk (**Fig. S3E**). These caspase cleavage events, then, could be responsible for the IR phenomenon.

However, we found a similar shift in IR distribution in AMO-1 cells treated with 15 nM Cfz (*n* = 27,386; median = 2.2) (**Fig. 3C**) despite much less cytoxicity (∼20%) than the 18 nM Cfz treatment in MM.1S. As caspase cleavage correlates with degree of cell death, it therefore appears unlikely that cytotoxicity alone is responsible for IR. Notably, an even smaller shift was observed in IR for MM.1S with 10 µM melphalan treatment, also at ∼20% cytotoxicity (*n* = 24,247; median = 0.44; *p* < 2.2E-16 for IR distribution MM.1S 18 nM Cfz vs 10 µM melphalan).

Instead, after melphalan the greatest ΔPSI shift occurred with alt. exon cassettes (single cassette median = 1.06, *n* = 12,267, coordinated cassette median = 1.75, *n* = 1,417, **Fig. 3D**). Even if we only consider statistically significant IR events (*p* < 0.05), the ΔPSI distributions for the drug responses remain distinct (**Fig. S4A-B**). **Fig. 3E** compares the GO enrichment of all significant ASEs induced by Cfz and by melphalan and shows divergent classes of genes alternatively spliced. Therefore, while melphalan also affects alternative splicing it appears to do so via a different mechanism than Cfz (24).

Intriguingly, in the case of AMO-1 treated with 15 nM Cfz, we noticed the ΔPSI shift for alternative splice sites (alt. acceptor median = 0.40, *n* = 9,620; alt. donor median = 0.61, *n* = 8,116, **Fig. 3C**, **Supplementary Table S3**) were decreased compared to MM.1S treated with 18 nM Cfz. Notably, this finding also correlates with the lesser degree of splicing factor phosphorylation (**Fig. 2**). These findings illustrate a more pronounced IR phenotype after PI than DNA damage, irrespective of the amount of cell death, while alternative exonic splice site determination may be a dose-response behavior.

### Exogenous expression of SRSF1 wildtype and RS-domain mutants do not significantly alter splicing patterns

Having shown that proteasome inhibition can lead to both robust splicing factor phosphorylation as well as widespread IR of pre-mRNA, we next considered whether these processes are causally linked or whether they instead occur via parallel mechanisms. To initially investigate this question, we considered SRSF1 (also known as SF2 or ASF), a well-characterized member of the SR family of splicing factors and a putative proto-oncogene (16, 25). All members of this family contain RNA recognition motifs (RRM) and arginine- and serine-rich domains (RS) (16). In general, SR proteins recognize *cis*-acting splice enhancers on pre-mRNA and work to promote splicing by initially recruiting the spliceosome to these intron-exon junctions (16). We found that SRSF1 demonstrates upregulated phosphorylation at sites in both the RS1 and RS2 domain when MM cells are treated with Cfz (**Fig. 2A, Supplementary Table S4**). The current model of SRSF1 function suggests that 1) SR-protein kinases (SRPK)-mediated phosphorylation of RS domain leads to translocation into the nucleus, 2) further hyperphosphorylation by CLK1 causes association with the U1 spliceosome, and 3) partial dephosphorylation is required for splicing catalysis (16, 26, 27).

To study the effects of SRSF1 phosphorylation in MM, we exogenously expressed a wildtype (SRSF1-WT), phosphomimetic (SRSF1-SD), or phosphodead (SRSF1-SA) variant in AMO-1 plasma cells. We assumed an all- or-none model of SR protein phosphorylation, where exogenous SRSF1 mutants have all 20 serines in the RS1 and RS2 domains replaced with either an aspartate (SD) or an alanine (SA). Exogenously expressed SRSF1 proteins are tagged with a C-terminal mCherry, nuclear localization signal (NLS) and 3x FLAG-peptide (**Fig. 4A**). It is known that phosphorylation of the RS1 domain is necessary for nuclear localization (28, 29); the attempted forced nuclear localization of the SA mutant was chosen to probe potential splicing-level effects of phospho-dead SRSF1 interacting with the spliceosome. Immunoblot confirms expression of exogenous SRSF1 constructs, which migrate higher than endogenous SRSF1 (**Fig. S4E)**, and demonstrates lower expression than the high-abundance endogenous protein. Epi-fluorescent images in **Fig. 4B** show the distribution of exogenous SRSF1-WT, SD, and SA mutants. Notably, most of WT and SD signal is localized to the nucleus, suggestive of functional protein product and consistent with expected biology. However, a much larger fraction of SA mutant is trapped in the cytosol despite NLS tagging. Consistent with prior work (30, 31), this finding suggests that phosphorylation of RS domains is a major requirement for entry into the nucleus.

**Figure 4.**
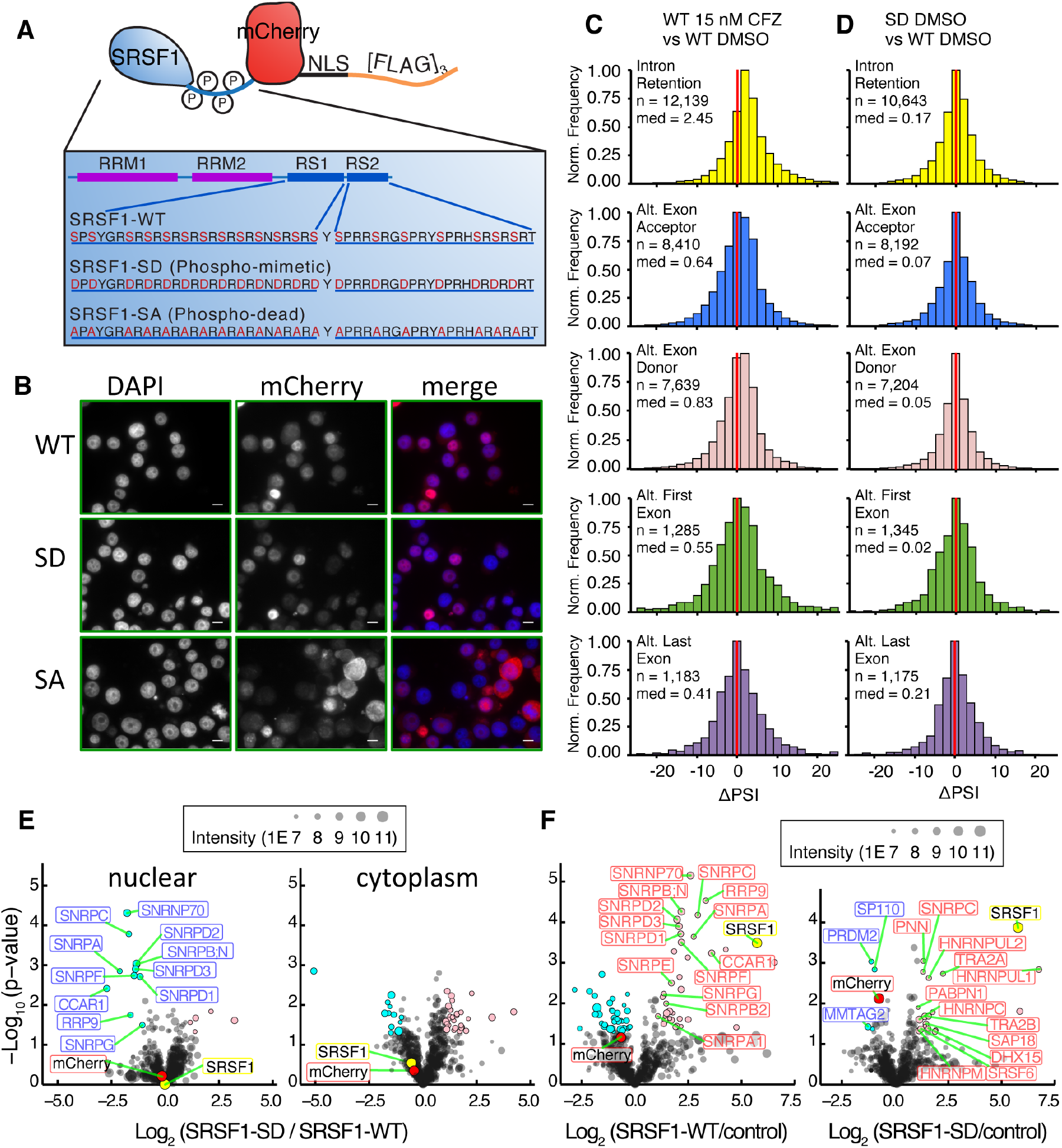
Modeling SRSF1 phosphorylation in MM drives interactome dynamics but not global splicing changes. **A.** Cartoon of protein architecture for exogenous SRSF1-NLS-mCherry-[FLAG]_3_. **B.** Epi-fluorescent imaging of DAPI stained AMO-1 expressing mCherry labeled SRSF1-WT (top), SRSF1-SD (middle), and SRSF1-SA (bottom). Scalebar represents 10 µm. **C and D.** Histograms of ΔPSI for IR, alt. exon donor, alt. exon acceptor, alt. first exon, alt. last exon ASEs when comparing differential splicing of AMO-1 expressing **C.** SRSF1-WT treated with 15 nM Cfz to DMSO and **D.** SD to WT. **E.** Volcano plots indicating differential interactors of SD compared to WT in both the nucleus and cytoplasm. **F.** Volcano plots of WT or SD when compared to control (NLS-mCherry-[FLAG]_3_) in AMO-1 nucleus reveal SD exclusion from spliceosome. Significant enriched proteins in pink, unenriched proteins in cyan (*p* < 0.05, ≥2-fold change). Circle size corresponds to summed LFQ intensities. mCherry ratio is red and SRSF1 ratio is yellow.

Upon JuncBASE analysis of poly-A RNA-seq data from DMSO-treated WT, SD, and SA construct (*n* = 3 for each), we saw remarkably few global differences in PSI as a function of modeled SRSF1 phosphorylation status (**Fig. 4D**). Notably, our results in **Fig. 1B** suggested that phosphorylation of multiple splicing factors, including other SRSF proteins, occurs simultaneously under Cfz-induced stress; we find that altered phosphorylation of SRSF1 alone may not carry any significant effects.

### SRSF1 RS-domain phosphomimetic mutant demonstrates weakened interaction with the spliceosome

Though we cannot draw a direct link between SRSF1 phosphorylation status and specific alternative splicing events, we further investigated the diverse biological roles of SRSF1. In addition to modulating pre-mRNA splicing, these include regulating nuclear export of spliced mRNAs and translational regulation in the cytosol via interaction with the ribosome (32-34). Using the 3x-FLAG tag on constructs we performed affinity purification mass spectrometry (AP-MS) with label-free quantitative proteomics vs. an mCherry-NLS-[FLAG]_3_ control. We specifically evaluated differential binding partners of SRSF1 as a function of phosphorylation status across both the nuclear and cytoplasmic compartments.

While clear differences were observed between the nuclear and cytosolic interactome for each construct, overall biological signatures based on GO analysis were surprisingly similar across WT, SD, and SA within each compartment (**Fig. S5B, D, F**). Notably, in the cytosol we found consistent interactions between both SRSF1-WT and SRSF1-SD with several RNA-binding proteins as well as components of the translational machinery. We do note one stark difference between WT and the phosphomimetic mutant in the nuclear fraction: the WT construct showed direct evidence of interaction with several small nuclear ribonucleoprotein (snRNP) polypeptides, core components of the U1-U2 spliceosome (**Fig. 4E**). Unexpectedly, these nuclear interactions were not enriched in the SD construct, which instead interacted with other splicing-related factors such as TRA2A, TRA2B, and PABPN (**Fig. 4F**). This interactome mapping may help refine the current model of SRSF1 biology, which suggests that hyperphosphorylation of RS domains leads to preferential integration with the U1 spliceosome (35, 36) and would explain the lack of change seen in global alternative splicing in the SD expressing cells.

### Proteasome inhibition of MM cells results in both stochastic intron retention and specific alternative exon usage

We next explored the splicing-level effects of 15 nM Cfz treatment on AMO-1 cells expressing the WT, SD, and SA constructs. Notably, in this setting cytotoxicity at 24 hr was <10% in 8 of 9 total replicates (**Fig. S4D**). Compared to DMSO-treated samples (**Fig. 4C**), Cfz again elicited a response consistent with that found in **Fig. 3C**: despite minimal cell death, we observed a clear shift in the median ΔPSI toward increased global IR (*n* = 12,139; median = 2.45, *p* < 2.2E-16 for one-sample Wilcoxon summed rank test). These findings in the absence of apoptosis underscore that caspase cleavage of splicing factors is unlikely to be a primary mechanism of IR after PI.

The combined RNA-seq dataset of all Cfz-treated samples vs. DMSO for these additional SRSF1 constructs were analyzed together (**Fig. 5A**) with JuncBASE (*n* = 24 replicates total across all AMO-1, including data in **Fig. 3C, 4C**). With this increased statistical power, we were able to identify *CNNM3*, which encodes a divalent metal cation transporter, as showing among the strongest signatures of IR across all events (FDR-corrected *p* = 0.032) (**Fig. S4G**). However, despite detecting *n* = 22,559 IR events by JuncBASE (**Fig. 5B, left**), very few individual transcripts (*n* = 43, including *CNNM3*) showed statistically significant (FDR-corrected *p* < 0.05) IR across replicates (**Fig. 5B, right**). This finding suggests that Cfz-induced IR may be a stochastic process, perhaps resulting from general interference with the splicing machinery without a coherent selection for specific transcripts.

**Figure 5.**
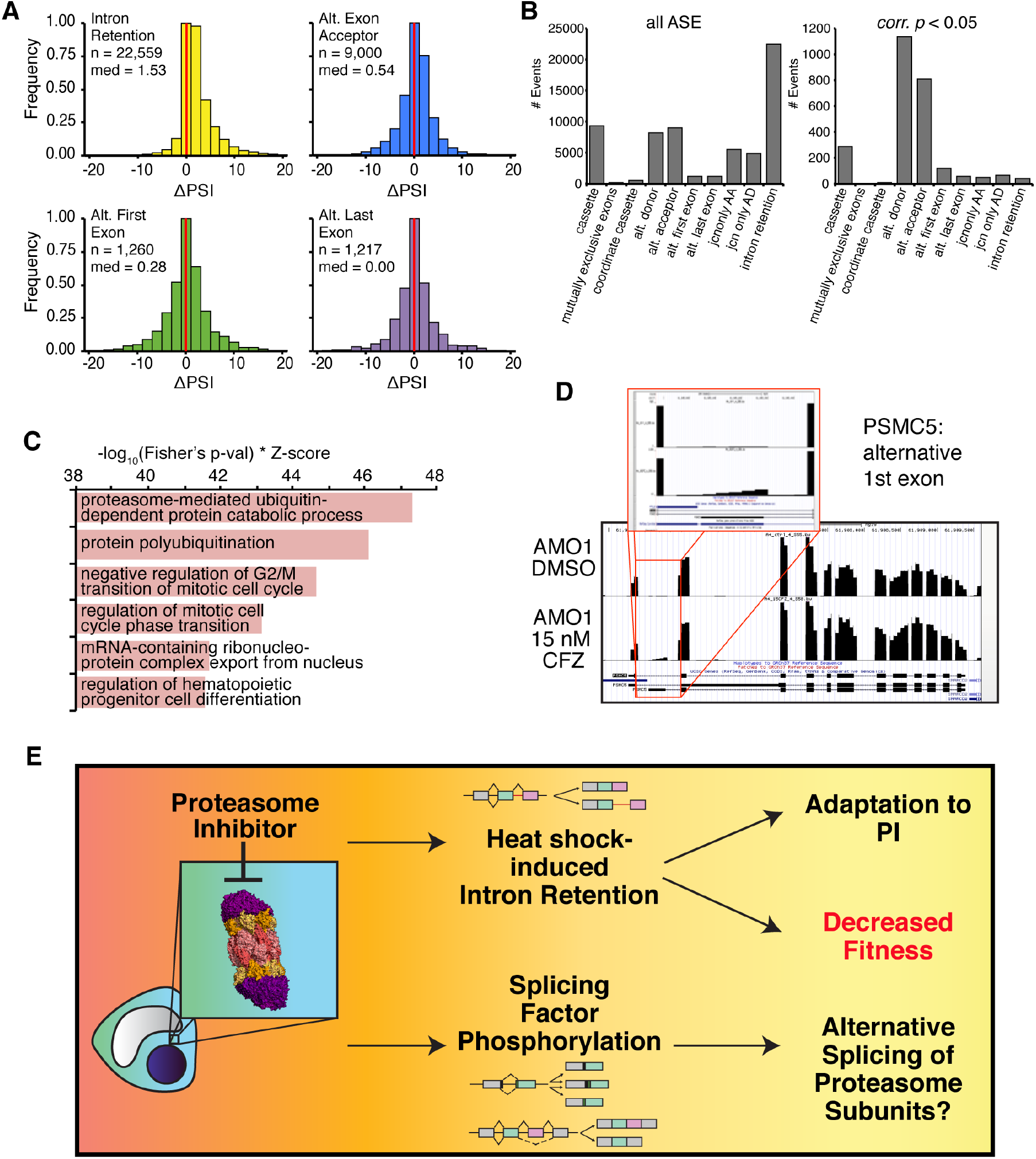
Combined SRSF1 constructs validate the splicing phenotype after Cfz. **A.** Histograms of ΔPSI in pooled analysis of parental AMO-1, AMO-1 SRSF1-WT, SD, and SA expressing cells treated with 15 nM Cfz compared to DMSO. **B.** Graph shows total number of events (n = 62,474) for each ASE type (left) and only the significant (FDR-corrected *p* < 0.05) events (n = 2,575) for each type (right). **C.** Top ranked GO enrichment terms of all genes involved in significant ASEs, regardless of type. **D.** Snapshot of UCSC Genome Browser Bar graph compares RNA-seq counts for the proteasomal subunit *PSMC5* between AMO-1 treated with 15 nM Cfz (bottom) and with DMSO (top). Inset displays sequencing counts showing alternative first exon. **E.** Model of new PI mechanism of action found in MM.

In contrast, alternative exon splice site usage (alternative exon donor (*n* = 1134) and alternative exon acceptor (*n* = 810)) emerged as the dominant type of alternative splicing when considering only statistically significant events (**Fig. 5B**). We investigated whether these consistently observed alternative splicing events may carry some biological relevance. Interestingly, GO enrichment analysis of all the genes undergoing significant alternative splicing after Cfz (n = 2,575 events total across all categories in **Fig. 5B, right**) revealed ‘proteasome-mediated ubiquitin-dependent protein catabolic process’ (*p* = 2.08E-16) and ‘protein polyubiquitination’ (*p* = 1.39E-13) as highly enriched (**Fig. 5C**). Notably, multiple proteasome subunits (*PSMA3/5/7, PSMB4/5, PSMC1/4/5, PSMD1-4, PSME2*), the protein homeostasis node p97 (*VCP*), and ubiquitin (*UBB, UBC*) all undergo some degree of alternative splicing with Cfz (example in **Fig. 5D**). These findings raise the possibility that alternative splicing may modulate the protein homeostasis machinery in response to therapeutic proteasome inhibition.

Taken together, our results offer a model for the effects of proteasome inhibition on the splicing machinery in myeloma (**Fig. 5E**). Upon therapeutic insult, the stress response induces phosphorylation of multiple splicing factors. Though the effect of this phosphorylation on specific splicing events remains unclear, these events may relate to specific alterations in exon usage based on known SRSF biochemistry. Our analysis of specific exon usage suggests that modification of the proteasome itself via alternative splicing may play a role in adaptation or resistance to proteasome inhibitor. In parallel, we observe a broad increase in the number of stochastically distributed IR events. These IR events, expected to reduce the number of functional protein products, may work to generally reduce proteotoxic stress and conserve cellular resources normally devoted to protein synthesis, thereby playing a role in adaptation to proteasome inhibition. Alternatively, the intron retention phenotype may indicate malfunction of the spliceosome, an essential process whose loss reduces tumor cell fitness. Interference with splicing may therefore be a previously unappreciated part of the PI mechanism of action.

### The spliceosome inhibitor E7107 is broadly potent versus MM cells and synergistic with proteasome inhibitor

Extending from this potential new mechanism of action of PIs as interfering with splicing, we further investigated the therapeutic potential of more dramatic spliceosome disruption in myeloma. For our preclinical studies we employed the tool compound E7107, a pladienolide B analog and direct inhibitor of the core U2 catalytic spliceosome component SF3B1 (15). This molecule has recently been described to induce extreme IR and strong cytotoxic effects versus models of myeloid malignancy, particularly those carrying mutations within splicing factors (37).

Using both qPCR validation of canonical IR events after SF3B1 inhibition (38) (**Fig. S6B**) as well as JuncBASE analysis of RNA-seq data (**Fig. 6A**), as expected we identified very significant IR after 6 h of 10 nM E7107 treatment in MM.1S cells (ΔPSI median = 13.79, *n* = 30,666). There was no noted cytotoxicity at this early time point (**Fig. S6A**). This finding supports the previous conclusion that splicing impairment, not apoptosis, induces IR. However, unlike the PI response, we also observed massive global loss of cassette exon splicing under E7107 (ΔPSI median = −16.6, *n* = 24,053, **Fig. S6D**). Furthermore, the number of significant (*p* < 0.05) IR events remained very high with E7107 (*n* = 7,171), unlike the apparently stochastic IR events seen with PI (**Fig. S6C**). Altogether, this suggests that PI-induced impairment of splicing is a partial interference of normal splicing operations, unlike the total abrogation of splicing seen with E7107. Underscoring the potential of splicing inhibition as a therapeutic strategy in MM, E7107 was extremely potent versus a panel of seven MM cell lines treated for 48 hr, with LC_50_’s ranging from <1 nM to 30 nM (**Fig. 6B**). In addition, a PI-resistant AMO-1 cell line (39) showed very similar sensitivity to E7107 as the parental line (**Fig. 6C**). This finding suggests the potential for clinical utility of splicing inhibition even in PI-refractory disease.

**Figure 6.**
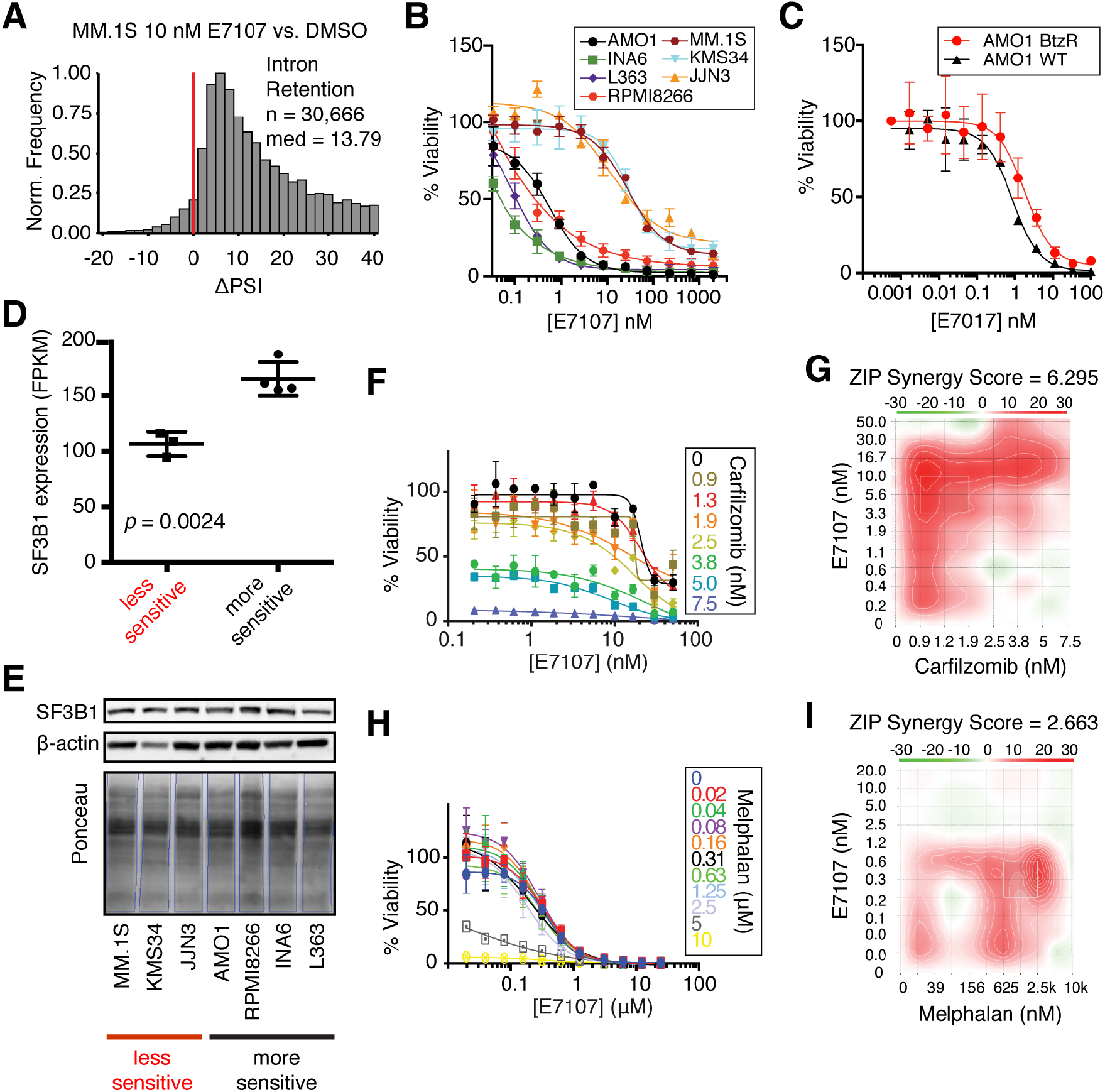
The catalytic spliceosome inhibitor E7107 induces IR and has potent anti-MM activity *in vitro*. **A.** ΔPSI histogram of IR events for MM.1S treated with 10 nM E7107 for 6 hr with respect to DMSO. Bin = 2 and red line at ΔPSI = 0. **B.** Cell viability curves compare a panel of 7 MM cell lines treated with E7107 for 48 hr (n = 4; mean +/- S.D.). **C.** Cell viability comparison of WT AMO-1 and PI-resistant AMO-1 cell line (BtzR) when treated with E7107 (n = 4; mean +/- S.D.) **D.** Evaluation of SF3B1 expression by RNA-seq (www.keatslab.org, mean +/- S.D.) and **E.** Western blot across more E7107- sensitive (AMO-1, INA6, L363, and RPMI8266) and less E7107-sensitive (MM.1S, KMS34, JJN3) cell lines. **F.** Cell viability curves of MM.1S combination therapy with E7107 and Cfz (n = 4; mean +/- S.D.) **G.** 2-D heatmap of ZIP synergy-scored landscape from Cfz and E7107 combination study. Red = synergy; green = antagonism; white = additive. Overall ZIP score of 6.295 suggests strong synergy. **H and I.** Same as **F.** and **G.** but for melphalan + E7107 combination. Overall ZIP synergy score of 2.663 denotes weaker synergy than with Cfz.

We noted that our MM cell line sensitivities appeared essentially bimodal, with one group of more sensitive lines with LC_50_’s of <1 nM and another slightly less sensitive group of cell lines with LC_50_ of 20-50 nM. In an attempt to identify potential determinants of this differential drug sensitivity, we examined publicly available RNA-seq data of baseline gene expression in MM cell lines (www.keatslab.org). We were intrigued to find that the more sensitive lines demonstrated significantly higher RNA expression of *SF3B1* (**Fig. 6D**). This outcome hinted that more sensitive cell lines may somehow be more “addicted” to SF3B1, leaving them more vulnerable to splicing inhibition, as well as revealing a potential biomarker that could be used for patient stratification. Unfortunately, this result was not confirmed at the protein level (**Fig. 6E**), suggesting that SF3B1 may undergo post-transcriptional regulation. We found no other candidates for markers of sensitivity or resistance to E7107 based on available DNA or RNA sequencing data from this limited cohort of cell lines.

We further explored the hypothesis that interfering with splicing via two different mechanisms may lead to synergistic MM cell death. Indeed, combination studies with Cfz and E7107 showed strong synergy across the dosing landscape based on ZIP synergy scoring (40) (**Fig. 6F-G**). In contrast, melphalan, which induced much less IR than PI (**Fig. 3B-C**), showed much weaker synergy in combination with E7107 (**Fig. 6H-I**). These findings support the approach of using splicing inhibitors in combination with PIs in MM treatment. Also, this result strengthens the hypothesis that splicing interference is a part of the PI mechanism of action.

### E7107 is a highly potent versus myeloma both *in vivo* and *ex vivo*

Based on this encouraging *in vitro* data, we then moved into a standard *in vivo* MM model of luciferase-labeled MM.1S cells implanted intravenously into NOD *scid* gamma (NSG) immunocompromised mice. These cells home to the murine bone marrow, partially recapitulating the tumor microenvironment in human disease (41). We found that E7107 was generally well-tolerated with no appreciable weight loss (**Fig. S7A**). At 3 mg/kg E7107 I.V., a relatively low dose compared to prior studies in other malignancies (37), we still found pronounced anti-MM effect after a brief 2 week treatment (**Fig. 7A-C**). This suppression of tumor translated into a significant survival benefit (*p* = 0.01, log-ranked test; *n* = 6 per arm).

**Figure 7.**
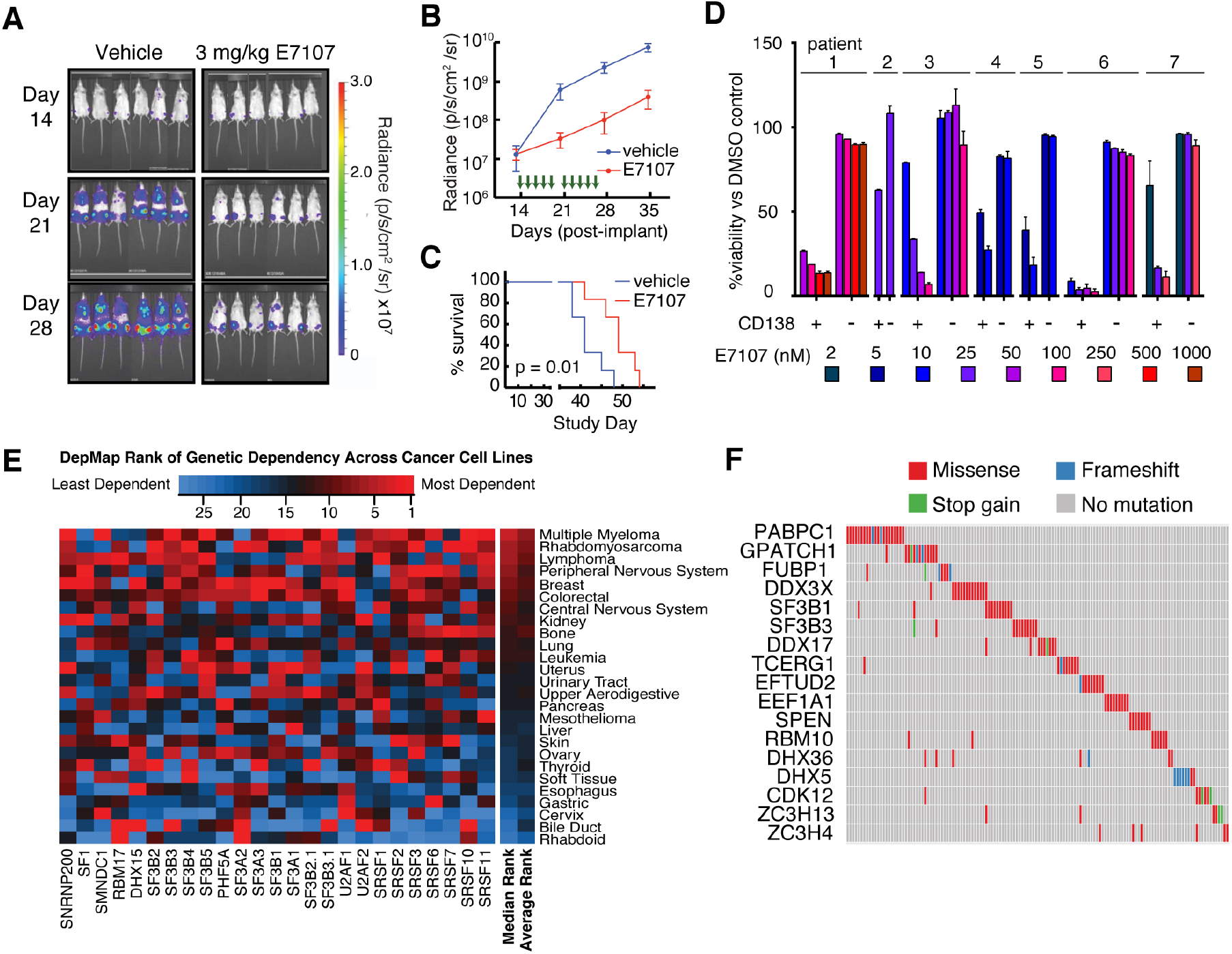
Inhibition of the spliceosome is a promising therapeutic strategy in myeloma. **A-B.** Bioluminescence imaging of luc-labeled MM.1S cells implanted in mice treated with either vehicle (left, n = 6) or 3 mg/kg E7107 (right, n = 6). Green arrows indicate days when drug was administered (14-18, 21-25). **C.** E7107 leads to significant improvement in murine survival (*p* = 0.01 by log-ranked test) **D.** Treatment of primary bone marrow aspirate samples from PI-refractory myeloma patients at various doses of E7107 for 24 hr shows significant cytotoxicity to CD138+ MM plasma cells at low-nM concentrations but minimal effects on other (CD138-) hematopoietic cells (n = 2 technical replicates; mean +/- S.D.). **E.** Heatmap of CRISPR-Cas9 essentiality screen data analysis in the Cancer Dependency Map (www.depmap.org; **Avana 18Q4** release) of core spliceosomal subunits among all tested tumor cell types. **F.** Analysis of MMRF CoMMpass data (research.themmrf.org; release **IA11**) summarizing mutations with possible functional effects in numerous splicing-related factors, as defined by Seiler *et al*. (46), within MM patient plasma cells.

We next turned to *ex vivo* evaluation versus primary patient samples. Fresh bone marrow mononuclear cells from seven PI-refractory MM patients were treated for 48 hr with varying doses of E7107. Based on flow cytometry analysis of CD138+ plasma cells (**Fig. S7B**), we found similar high sensitivity of patient tumor cells to E7107 as found in cell lines, with estimated LC_50_’s in the low-nM range (**Fig. 7D**). Notably, non-plasma cell bone marrow mononuclear cells (CD138- fraction) showed remarkably little cytotoxicity at these same doses, supporting a potential therapeutic index for splicing inhibitors in MM.

### Functional genomics and whole exome sequencing suggests clinical applications of splicing inhibition in MM

Analysis of CRISPR essentiality screen data in the Cancer Dependency Map (www.depmap.org; Avana library public 18Q4 (42)), across over 400 cancer cell lines, demonstrated that myeloma has among the strongest genetic dependencies on the target of E7107, *SF3B1* (**Fig. S7C**). This genetic ablation data further supports the ability to pharmacologically eliminate MM tumor cells via splicing inhibition while sparing normal cells. We further extended this analysis to other core components of the U1-U2 spliceosome found to be “common essential” genes per DepMap (43). By aggregating DepMap rankings, we found that MM cell lines are the most sensitive tumor cell type to genetic ablation of these central snRNP protein components, necessary for association with pre-mRNA and splicing catalysis (**Fig. 7E**). Compared to the essential subunits of the 20S proteasome (including the direct PI target *PSMB5*) (**Fig. S7D**), we surprisingly found more favorable genetic evidence for targeting the spliceosome in MM than the proteasome.

Furthermore, a recent study validated the sulfonamide indisulam as an inhibitor of splicing via targeted degradation of RBM39, another component of the spliceosome with high homology to U2AF2 (44). In this work, hematopoeitic malignancy cell lines were broadly more sensitive to indisulam than solid tumor cell lines. We confirmed cytotoxicity of indisulam versus a panel of MM cell lines (**Fig. S7E**), though LC_50_’s (0.3 - >20 µM) were much higher than those for the SF3B1 inhibitor E7107. In DepMap data, MM was again among the more sensitive tumor type to RBM39 ablation (**Fig. S7F**). Indisulam may therefore represent another approach to targeting the spliceosome in this disease, though given lower potency the potential for clinical translation is less clear.

We next took advantage of genomic and transcriptomic data from isolated malignant plasma cells from newly-diagnosed MM patients in the Multiple Myeloma Research Foundation CoMMpass study (research.themmrf.org; version IA11). First evaluating gene expression data, we intriguingly found significantly decreased progression-free survival among patients in the top quartile of *SRSF1* expression versus those in the bottom quartile (*p* = 0.0081 by log-ranked test) and a trend toward similarly decreased overall survival for patients in the top vs. bottom quartile of *SF3B1* expression (*p* = 0.087) (**Fig. S7G**). These results raise the possibility of poorer outcomes in patients whose disease is more dependent on the spliceosome.

However, we note that both E7107 (37) and the recently described splicing inhibitor H3B-8800 (38) have both been shown to have the greatest potency versus hematopoietic malignancies carrying mutations in splicing factors such as *SF3B1*, *SRSF2*, *U2AF2*, and *ZRSR2* (45). These mutations are seen frequently in myelodysplastic syndromes (MDS), acute myeloid leukemia, and chronic lymphocytic leukemia, appearing in up to 50% of MDS patients (45). We therefore examined exome sequencing data available in CoMMpass and found that 28.0% of MM patients (268 of 956) were found to carry missense mutations within at least one of 119 splicing-associated factors recently proposed to be most relevant to tumorigenesis across a survey of the The Cancer Genome Atlas (**Fig. 7F, Supplementary Table S6**) (46). While only a small minority of these identified mutations has been functionally validated to affect splicing, the most common single mutation was at the known “hotspot” *SF3B1* K666T, found in three patients. Variant allele frequencies for these expected heterozygous mutations were 42%, 35%, and 22%, suggestive of a prominent subclonal fraction of the tumor cell population. Among well-characterized genes, mutations were found in *SF3B1* (*n* = 10 patients, including K666T mutations), *SRSF2* (*n* = 2), *U2AF1* (*n* = 4), and *ZRSR2* (*n* = 1). Unfortunately we were unable to obtain rare primary patient samples containing mutations in these genes, and no myeloma cell lines are known to carry hotspot mutations in these well-characterized splicing factors (www.keatslab.org). While our data suggest that spliceosome inhibition should be considered a therapeutic option for MM patients of any genotype, recent work in other malignancies (37, 38) supports the potential for particular benefit in the subset of patients carrying pathogenic splicing factor mutations.

## DISCUSSION

Our results demonstrate that PI therapy in myeloma leads to both specific alterations in splice site usage and broad-scale interference with spliceosome function. This observation, initially generated through unbiased phosphoproteomics, led us to explore the spliceosome itself as a MM vulnerability. Our preclinical evaluation and analysis of functional genomics and exome sequencing data further reinforced the spliceosome as a therapeutic target in MM.

These results raise a number of intriguing questions. From a mechanistic perspective, prior work examining SR phosphorylation after cellular perturbation using Western blotting did not consistently show a broad hyperphosphorylation signature (47-49). Our results therefore illustrate the utility of unbiased phosphoproteomics to elucidate cancer drug response. Recent work suggests that additional kinases beyond the well-characterized SRPKs and CLKs may be involved in SR phosphorylation (50, 51). However, in the context of drug-induced stress in cancer, the mechanism that leads to coordinated, upregulated phosphorylation across multiple splicing factors, whether via kinase activation or phosphatase inhibition, will be an important topic for future investigation.

We also found a correlation between PI-induced stress and both SR factor phosphorylation and the degree of alternative exon selection. This finding appears consistent with prior studies (16) suggesting that SR factors play a role in exon selection and their activity can be modulated by phosphorylation. Why these effects specifically lead to alternative splicing of protein homeostasis genes remains to be investigated. In contrast, we found IR to be largely independent of the degree of PI-induced stress. This result suggests that the IR phenotype is mediated by a different mechanism and uncoupled from SR factor phosphorylation.

In attempting to model the relationship between SR phosphorylation and splicing, we recognize our phosphomimetic construct likely does not fully recapitulate the complex phosphorylation biology of SRSF1 within cells (27), nor does it fully match the level of expression of endogenous SRSF1. In general, causal links have been noted between SRSF1 phosphorylation and splicing in single transcript, *in vitro* systems (34, 52, 53), but isolating global effects of SR phosphorylation on splicing within cells have remained elusive. Furthermore, using genetic approaches we cannot readily model phosphorylation changes on multiple SR proteins simultaneously, which may be necessary to elicit broader phenotypic effects. Despite these limitations, however, our WT expression studies provide a landscape of the SRSF1 cytosolic and nuclear interactome, which may inform future studies of SR protein biology in myeloma and other systems.

This PI-induced interference with normal splicing even at minimal cytotoxicity, much greater than that found with melphalan, may relate to the activation of the heat shock response. We found prominent heat shock chaperone induction even under a non-cytotoxic dose of the PI bortezomib in MM.1S cells (10). As previously shown in non-cancer cells, heat shock alone, without cell death, can lead to significant intron retention (23). One hypothesis is that this broad-scale inhibition of splicing acts in a similar fashion to translational inhibition after drug-induced stress: a way to conserve cellular resources and focus on only producing genes required for survival and the stress response. However, as described in our model of **Fig. 5E**, and evidenced by our mRNA-seq data after E7107 treatment (**Fig. 6A**), another possible result of widespread intron retention and downstream loss of normal protein production is significant decrease in cellular fitness and ultimately, cell death. There may be a quantitative threshold effect between these two outcomes that remains to be elucidated.

Here, we propose that the loss-of-fitness modality of drug-induced IR constitutes a previously unexplored mechanism of action of PIs. We further performed a preclinical evaluation of splicing inhibition in myeloma using E7107, finding potent anti-myeloma effects *in vitro*, *in vivo*, and *ex vivo* versus primary patient samples. From a therapeutic perspective, one of the major questions is the potential toxicity of targeting such an essential process as the catalytic spliceosome. However, our analysis of genetic dependency data and our *ex vivo* data with E7107 clearly demonstrates the potential to target core spliceosome subunits in MM while largely sparing normal cells. In fact, based on this analysis the spliceosome appears to be an even more promising target than the clinically-validated approach of targeting essential subunits of the proteasome. Furthermore, presumed efficacious doses (based on measured blood concentrations in the nM range) of E7107 were largely well tolerated in a Phase I clinical trial (54). While this molecule is no longer in clinical development, it is thought that E7107 visual toxicity was molecule-specific and is not a function of targeting the spliceosome in general (15). Our genomic analysis suggests that mutations in splicing factors are found in a substantial fraction of MM patients. Newer generations of splicing inhibitors are currently in clinical trials for other hematologic malignancies (38) (NCT02841540) and may be of particular benefit for these patients. Our results support clinical investigation of these compounds in MM either alone or to enhance PI efficacy as combination therapy.

## METHODS

### Cell culture

All cell lines were grown in suspension at 37°C, 5% CO2 in complete media: RPMI 1640 medium (Gibco, 22400105, UCSF CCFAE002), supplemented with 10% FBS (Atlanta Biologicals, S11150) for proteomics experiments and Benchmark FBS (Gemini Bio-products, 100-106) for drug viability experiments and 1% penicillin-streptomycin (UCSF, CCFGK003). INA6 cell media was supplemented with 90 ng/mL recombinant human IL-6 (ProSpec Bio, CYT-213).

#### Drug cytotoxicity assay

For dose-response cell toxicity assays, 1E+3 myeloma cells were seeded per well in 384 well plates (Corning) using the Multidrop Combi (Thermo Fisher) and incubated for 24 hr. In monotherapy cytotoxicity assays, cells were treated with drug or DMSO and incubated for 48 hr, while cells were further incubated with E7107 (H3) for an additional 24 hr in E7107 dual therapy combination assays. Carfilzomib (SelleckChem, S2853-50 mg), melphalan (Sigma, S2853-50 mg), and E7107 (H3 Biomedicine, CAS:630100-90-2), and KH-CB19 (sc-362756) were solubilized in DMSO at 10 mM.

All cell viability was determined with Cell-Titer Glo reagent (Promega, G7573) using a Glomax Explorer (Promega) luminescence plate reader. For the drug titration cytotoxicity assays, measurements were performed in quadruplicate, while measurements were performed in triplicate in all other assays, and viabilities are reported as mean (+/- S.D.) ratio normalized to DMSO-treated controls or measurements at 0 hr. For ZIP synergy calculations, normalized viability data was submitted to SynergyFinder web application (40).

#### Drug dosing for proteomics and RNA-seq experiments

Proteomic/phosphoproteomic/RNA-seq experiments were performed at a cell density of 1E6 cells/mL. For timecourse studies, ∼20E6 cells were grown in complete media for each timepoint (0, 8, 16, and 24 hr), whereas for single-timepoint experiments, 15-20E + 6 cells in light SILAC media were treated with drug compound and cells in heavy SILAC media (L-Lysine-^13^C_6_,^15^N_2_, L-Arginine-^13^C_6_,^15^N_4_ (Cambridge Isotope, CNLM-291-H-1, CNLM-539-H-1) were treated with DMSO for 24 hr. 1-3E+6 cells were set aside for RNA-seq. Cells were washed in PBS and cell pellets were frozen in liquid nitrogen (LN2) and stored in −80°C. 1 biological replicate for the timecourse experiment, 2 biological replicates for each single-timepoint condition (with a third only for RNA-seq), and 3 biological replicates for all AP-MS were gathered and analyzed.

#### Cloning and lentiviral transduction

SRSF1 and mCherry genes, along with 3X FLAG sequences and nuclear localization signal (NLS) were cloned into pLV-416G second generation lentiviral plasmid (UCSF HMTB) by Gibson Assembly. SRSF1 constructs were transfected into Lenti-X 293 T(Takara Bioscience, 632180) packaging cells with Gag-Pol expressing pCMV-dR8.91 (Addgene, Plasmid#2221) and VSV-G envelope expressing pMD2.G (Addgene, Plasmid#12259) plasmids. Viral particles were harvested, concentrated with Lenti-X concentrator (Takara Bioscience, 631231) and viral titers were incubated with AMO-1 cells. Positively transduced cells were selected with selection drug, G418 (VWR, 970-3-058), for several passages, then by mCherry expression with Fluorescence Activated Cell Sorting (FACS, Sony SH800). Protocol details are found in Supplementary Information.

#### Phosphoproteomic peptide preparation

Frozen pellets of ∼15-20E6 cells were lysed in 8 M urea, 0.1 M Tris pH 8.0, 150 mM NaCl and 1X HALT phosphatase/protease inhibitor cocktail (Pierce, 78442) for timecourse experiments or 8 M Guanadine-Cl (Gdn, Chem Impex Intl., 00152- 1KG), 0.1 M Tris pH 8.5, 10 mM tris(2-carboxyethyl)phosphine (TCEP, Pierce, 20491), 40 mM 2-chloroacetamide (2-CAA, Sigma, 22790-250G-F), 1X HALT for SILAC samples and lysed with probe sonicator (BRANSONIC). In the case of single-timepoint SILAC samples, equal part light and heavy labeled lysate samples were combined (∼ 2.5–3 mg total). Lysate is diluted with 0.1 M Tris pH 8.0 to a final concentration of 1.3 M Gdn or urea. Proteome is digested with 1:100 dilution of trypsin overnight for 22-24 hr at room temperature. Peptides are extracted with SEP-PAK C18 cartridges (WATERS). For single-timepoint SILAC samples, ∼100 µg of eluted peptides were dried and analyzed separately by LC- MS/MS as unenriched “global proteomics.” Remainder of eluate was diluted 3-4 fold with water, lyophilized, then resuspended in 80% ACN, 0.1% TFA and enriched on FeCl3 charged NTA-agarose beads sitting atop a C18 matrix in a stage-tip platform (Nest). Eluted phosphopeptides are dried and stored at −80°C.

#### Affinity Purification

For each replicate, frozen cell pellets were gently lysed on ice with 200 µl hypotonic lysis buffer (20 mM Tris (pH 7.4@4°C), 10 mM KCl, 0.1 mM EDTA (Fisher, BP120-500), 0.5% NP-40 alternative (EMD, 492016-100ML), 1 mM DTT (Gold Biotech, DTT50), 1 mM PMSF (RPI, P20270-1.0), 1x HALT protease/phosphatase inhibitor cocktail (Pierce, 78442), 300mM Sucrose, 0.03 U/mL aprotinin (RPI, A20550-0.001)), underwent 3 X freeze-thaw cycles, and clarified with 5 passes through an 18-gauge syringe needle. Lysate was centrifuged at 5,000 rcf, 4°C for 10 min and supernatant was reserved as cytoplasmic fraction, while nuclear fraction was washed and resuspended in 60 µl of 20 mM HEPES (pH 7.9), 420 mM NaCl, 25% glycerol, 1 mM EDTA, 1 mM DTT, 1 mM PMSF, 0.03 U/mL aprotinin, 1x protease/phosphatase inhibitor cocktail (HALT), 25 U Benzonase/mL and clarified with 10 passes through 18- gauge syringe needle. Both fractions were adjusted to 50 mM Tris pH 7.4, 150 mM NaCl, 1mM EDTA (binding buffer) and combined with M2 anti-FLAG magnetic beads (Sigma, M8823). Bound lysate was washed with binding buffer + 0.05% NP-40, then binding buffer, then twice with 20 mM Tris pH 8.0, 2 mM CaCl2. Proteins are denatured and cystines are reduced and alkylated with 6 M Gdn, 40 mM 2-CAA, 5 mM TCEP, 100 mM Tris pH 8.0, then trypsinized on-bead with ∼0.75 µg trypsin/ sample, ∼ 20 h at 37°C, and peptides were desalted with homemade C18 stagetips and dried and stored at −80°C.

#### LC-MS/MS

∼1 µg peptides were analyzed for each sample by “shotgun-“ LC-MS/MS on a Dionex Ultimate 3000 RSLCnano with 15 cm Acclaim PEPMAP C18 (Thermo, 164534) reverse phase column and Thermo Q-Exactive plus mass spectometer. Samples were analyzed with either a 3h 15 min non-linear gradient or a 1h 23 min linear gradient from 2.4% acetonitrile (ACN, Sigma, 34998-4L), 0.1% FA to 32% ACN. Experiment specific LC-MS/MS settings are listed in Supplementary Information.

#### Proteomic data analysis and quantification

Initial timecourse unlabeled phosphoproteomics data were processed together on Maxquant v1.5.1.2 (55) and searched against the human proteome (Uniprot downloaded 2014/12/3, with 89,706 entries). All AP-MS samples were processed together with similar settings. All SILAC samples (phospho-and unenriched peptides) were processed together with similar settings. SILAC quantification for global proteomics at the protein level requires 1 minimum razor or unique peptide. A one-sample T-test was applied to the log-2 transform of the normalized SILAC-labeled peptide ratios (heavy:light) for single-timepoint analysis, while for AP-MS data, two-sample T-test was applied to the log-2 transform of the median-normalized MaxQuant label-free quantification (LFQ) values of protein groups. The number of total entries (phosphosites, protein groups, significance is *p* < 0.05, |t-test difference| ≥ 1), along with correlation statistics between replicates, are summarized in **Supplementary Table S4** and shown in **Fig. S3B**. See Supplementary Information for specific search and analysis settings.

#### RNA-seq library preparation

RNA was extracted from frozen cell pellets with RNeasy Mini-prep kit (Qiagen, 74104). For timecourse experiments, cDNA library of expression transcripts was carried out with RNA Hyper Prep kit with RiboErase (Kapa, KK8560) to enhance transcript reads above ribosomal reads, while single-timepoint experiments assessing splicing required mRNA enrichment with magnetic mRNA Isolation kit poly-dT beads (NEB), then RNA Hyper Prep kit (Kapa, KK8540) for cDNA construction of 200-400 bp library with Illumina platform TruSeq indexed adaptors (**Supplementary Table S1**). RNA and DNA quantified at all steps by Nanodrop (Thermo Scientific) and cDNA library size and quality were evaluated on a Bioanalyzer 2100 (Agilent) with High Sensitivity DNA Kit (Agilent, 5067- 4626), before being submitted for next generation sequencing on a HiSeq4000 (Illumina) at the UCSF Center for Advanced Technologies core facility.

#### JuncBASE alternative splicing analysis

Alternative splicing events were identified and quantified with JuncBASE v1.2- beta using default parameters (21). Intron-exon junction database was created from hg19 annotations. T-test was used to compare number of inclusion and exclusion reads and p-values were adjusted with Benjamini-Hochberg correction. For ΔPSI histograms in Fig. **3B-C, 4C-D, and 5A**, JuncBASE output included a subset of alternative splice events with median PSI = 0.00 in both conditions or median PSI = 100.00 in both conditions, resulting in ΔPSI = 0.00. These events were manually removed for downstream analyses. Histograms and splicing statistics were determined with statistical computing program R (v3.5.1) and a summary is listed in **Supplementary Table S3**.

#### Gene Ontology enrichment analysis

Gene Ontology (GO) enrichment analysis of upregulated phosphosites and enriched SRSF1 interactors was performed in STRING (v10.5, https://string-db.org/) (56), searching against a background of all quantified protein entries. Enrichment analysis of all significantly alternative spliced genes (raw *p* < 0.05) was performed using web-based enrichment analysis tool, Enrichr (http://amp.pharm.mssm.edu/Enrichr/) (57). Reported combined score is calculated by multiplying the natural log transform of the *p*-value with the Fisher’s exact test of expected rank deviation (Z-score). Functional GO analysis is limited to biological processes and compiled in **Supplementary Table S5**.

#### Xenograft mouse model and *in vivo* luminescence imaging

1E6 MM.1S-luc cells, stably expressing luciferase, were transplanted via tail vein injection into 12 NOD.Cg-*Prkdc^scid^ Il2rg^tm1Wjl^*/SzJ (NSG) mice from The Jackson Laboratory (cat# 005557). All the mice were female, 6-8 wks old at start of studies, and typically weigh 20-25 g. NSG mice were handled with aseptic techniques and housed in pathogen free environments at the UCSF Laboratory Animal Research Center. All mouse studies were performed according to UCSF Institutional Animal Care and Use Committee-approved protocols. Tumor burden was assessed through weekly bioluminescent imaging in the UCSF preclinical therapeutic core on a Xenogen In Vivo Imaging System (IVIS), beginning 13 days after implantation, which is the same day as treatment initiation. Tumor implanted humanized mice were randomized and sorted into control arm and treatment arm, 6 mice/arm. Mice were treated for two weeks (five days on, two days off) with vehicle or 3 mg/kg E7107, formulated in vehicle (10% Ethanol, 5% Tween-80, QS with Saline) and administered by continuous subcutaneous infusion. Mice were kept and observed until survival endpoint; final timepoint was 54 days after MM.1S transplant. Acquired luciferase intensities were quantified with Living Image Software (PerkinElmer) in units of radiance (photons/s/cm2/sr). Kaplan-meier survival curves along with log-ranked test to determine significance were calculated in GraphPad Prism 6 software.

#### Patient bone marrow aspirate, CD138 labeling and flow cytometry analysis

Fresh de-identified primary multiple myeloma patient bone marrow (BM) samples were obtained from the UCSF Hematologic Malignancies Tissue Bank in accordance with the UCSF Committee on Human Research-approved protocols and the Declaration of Helsinki. Bone marrow mononuclear cells were isolated by density gradient centrifugation with Histopaque-1077 (Sigma Aldrich), and washed with 10 mL D-PBS 3 times. Mononuclear cells were resuspended in a small volume (∼1.5 mL) of media (RPMI1640, 10% FBS, 1% penicillin/streptomycin, 2 mM glutamine) and incubated at 37°C, 5% CO2 for 15 min. Isolated mononuclear cells from multiple myeloma patient bone marrow were adjusted to 2E5 cells/well in a 96 well plate. Cells were stimulated with 50 ng/ml recombinant human IL-6 (ProsPec) for 17 hr before treatment with E7107 or DMSO for 24 hr. Cells were then stained with 10 µL Alexa-Fluor 647 mouse anti-human CD138 antibody (BD Pharmingen, cat# 562097; RRID:AB_10895974) or Alexa-Fluor 647 IgG κ isotype (BD Pharmingen, cat# 557714; RRID:AB_396823) control and 2 µL SyTOX Green (Thermo, S34860) per 1 mL FACS buffer (D-PBS, 5% FBS). Resuspended cells are characterized with a CytoFLEX fluorescence cytometer (BD).

#### Statistical Analyses

All data are presented as mean +/- standard deviation, unless otherwise stated. Statistical significance in proteomics comparisons was determined by Student’s *t*- test: One-sample *t*-test with null hypothesis that log2-transform of the normalized SILAC ratio = 0, or a two-sample *T*-test with null hypothesis that the difference in log2-transform of the intensities is equal to. A *p* < 0.05 is considered statistically significant. For all Kaplan-Meier survival analysis, log-ranked test was used to determine statistical significance.

#### Data Availability

The mass spectrometry proteomics data and MaxQuant analysis results have been deposited to the ProteomeXchange Consortium via the PRIDE (58) partner repository with the dataset identifier PXD012172. Datasets are private during review; reviewers may access datasets with following credentials:

Username

Password: Xxe5zQtD

Raw RNA-seq data, processed analysis files, and JuncBASE results may be downloaded from the Gene Expression Omnibus, GEO (https://www.ncbi.nlm.nih.gov/geo/) with the accession number: GSE124510. Reviewers may access this data during review with the token: ifwxqgsgptcbtgz

## Acknowledgements

We thank Dr. Silvia Buonamici at H3 Biomedicine for providing E7107 and insightful discussions and Jacob Runyan for assisting in quality control of sequencing data. We thank Drs. Renate Burger and Christoph Driessen for providing INA-6 and AMO-1 parental and bortezomib resistant cell lines, respectively. We also thank the lab of James Wells for use of Zeiss Z1 Observer microscope, Center for Advanced Technology at UCSF for HiSeq sequencing, Dr. Jane Gordon and the Laboratory for Cell Analysis at Helen Diller Family Comprehensive Cancer Center for use and assistance of Sony SH800, and Dr. Danielle Swaney for discussion of AP-MS. This work was supported by NIH/NCI P30CA083103 (Cancer Center Support Grant, supporting UCSF Preclinical Therapeutic Core facility managed by B.C.H.), NIH/NHGRI T32HG008345 (to A.M.T.), the Damon Runyon Cancer Research Foundation Dale Frey Breakthrough Award (DFS 14-15), NIH/NCI K08CA184116, NIH/NIGMS DP2OD022552, and the UCSF Stephen and Nancy Grand Multiple Myeloma Translational Initiative (to A.P.W.) and NIH/NCI R01CA226851 (to A.N.B. and A.P.W.).

## Author Contributions

H.H.H., I.F., C.L., P.B., M.C.M., M.M., and A.P.W. performed experiments and analyzed experimental data. A.M.T., Y-H.L., and A.B. analyzed transcriptomic and genomic data. J.M., P.P., and B.C.H. performed *in vivo* studies. T.G.M., J.L.W., S.W.W., and N.S. consented patients and obtained primary specimens. H.H.H. and A.P.W. wrote the manuscript with input from all authors.

