## Supplementary Information for "Proteasome inhibitor-induced modulation reveals the spliceosome as a specific therapeutic vulnerability in multiple myeloma"

**This document includes:**

Supplementary Figures S1 – S7
Supplementary Tables S1 – S3
Supplementary Methods and References

**The following supplementary files are available as separate .xlsx files:**

Supplementary tables S4 – S6 (Excel Files)

***Corresponding Author Contact information:**
Arun P. Wiita, MD, PhD

UCSF Dept. of Laboratory Medicine
185 Berry St., Ste. 290
San Francisco, CA 94107

**List of Supplementary Items:**

**Supplementary Figure S1:** Time-course of MM cell phosphorylation after Cfz treatment.

**Supplementary Figure S2:** Protein abundance and gene expression response to drug perturbation

**Supplementary Figure S3:** Figure S3. Characterization of myeloma response to Cfz and melphalan.

**Supplementary Figure S4:** Cfz-induced splicing alterations across parental MM cells and SRSF constructs.

**Supplementary Figure S5:** The phosphorylation-dependent interactome of SRSF1.

**Supplementary Figure S6:** E7107 cell toxicity, functional splicing assay, and splicing statistics.

**Supplementary Figure S7:** Preclinical and clinical relevance of targeting the spliceosome in myeloma.

**Supplementary Table S1** Oligo sequences for cloning, qPCR, and RNA-seq cDNA library.

**Supplementary Table S2** Description of SRSF1 lentiviral expression constructs.

**Supplementary Table S3** Distribution statistics of ΔPSI calculated in R. Statistics for individual event types and all event types are listed for each comparative condition.

**Supplementary Table S4:** excel file with tabs listing comparative analysis (T-test *p*-value and $log_2$-difference) of phospho- and global proteomics

**Supplementary Table S5:** excel file with tabs summarizing bioinformatics GO enrichment analysis and KSEA kinase scores.

**Supplementary Table S6**: excel file listing splicing gene mutations and variant allele frequencies found in MM patients from CoMMpass dataset.

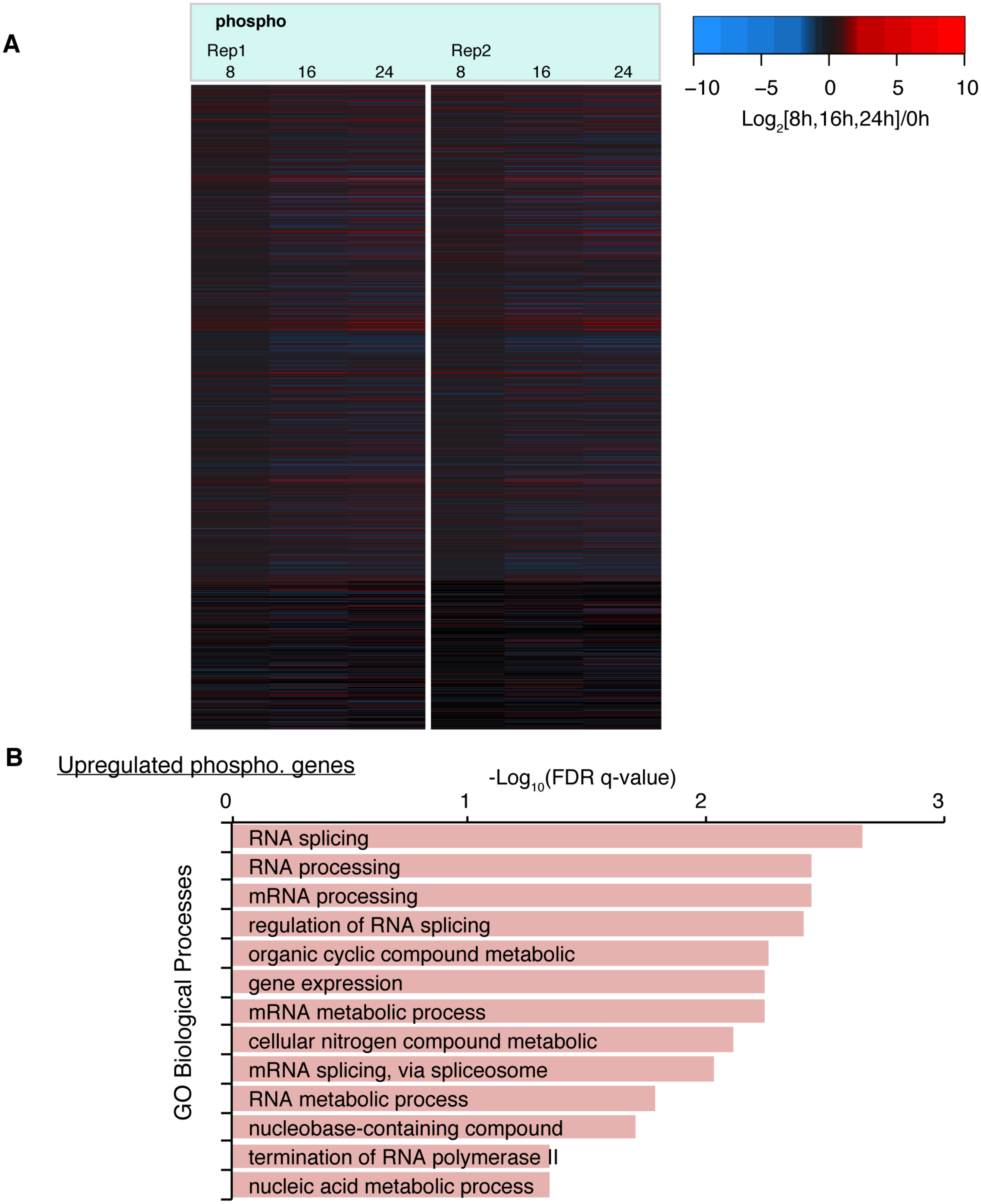

**Figure S1. Time-course of MM cell phosphorylation after Cfz treatment. A.** Heatmap of all 5791 quantified phosphosites, $log_2$-transformed label-free quantification (LFQ) intensity ratios (relative to 0 hr at 8, 16, 24 hr) for 2 technical replicates of MM.1S cells treated with 30 nM Cfz over a 24 hr time course. **B.** Top ranked (FDR *q*-value) GO enrichment terms for genes with increased phosphorylation and relatively unchanged transcript levels (**Fig. 1 B**) over 24 hr.

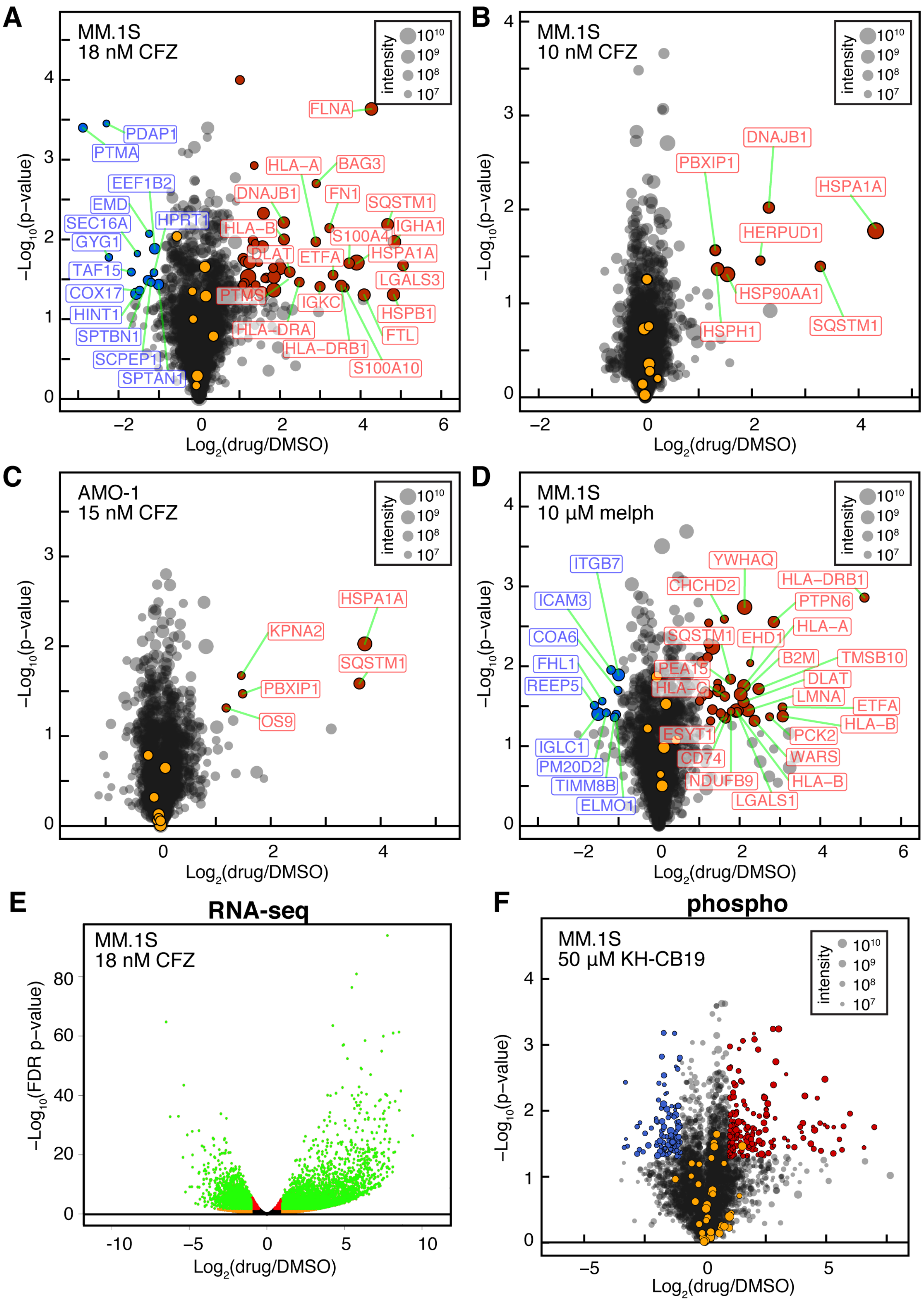
A
MM.1S
18 nM CFZ
intensity
FLNA
PDAP1
PTMA
EEF1B2
EMD
SEC16A
GYG1
HPRT1
TAF15
COX17
HINT1
SPTBN1
SCPEP1
SPTAN1
HLA−A
BAG3
DNAJB1
FN1
SQSTM1
HLA−B
IGHA1
S100A4
DLAT
ETFA
HSPA1A
LGALS3
PTMS
IGKC
HSPB1
HLA−DRA
FTL
HLA−DRB1
S100A10
Log2(drug/DMSO)
−Log10(p−value)
B
MM.1S
10 nM CFZ
DNAJB1
PBXIP1
HSPA1A
HERPUD1
HSP90AA1
HSPH1
SQSTM1
C
AMO-1
15 nM CFZ
HSPA1A
KPNA2
SQSTM1
PBXIP1
OS9
D
MM.1S
10 µM melph
ITGB7
YWHAQ
CHCHD2
HLA−DRB1
ICAM3
PTPN6
COA6
SQSTM1
EHD1
HLA−A
FHL1
PEA15
B2M
TMSB10
REEP5
HLA−C
DLAT
LMNA
ETFA
HLA−B
ESYT1
PCK2
IGLC1
CD74
WARS
PM20D2
TIMM8B
NDUFB9
HLA−B
ELMO1
LGALS1
E
RNA-seq
MM.1S
18 nM CFZ
−Log10(FDR p−value)
F
phospho
MM.1S
50 µM KH-CB19

**Figure S2. Protein abundance and gene expression response to drug perturbation.** Volcano plots showing $\log_2$ transformed ratios of single time-point SILAC-based LC-MS/MS protein intensities of MM cells treated with **A.** 18 nM Cfz, **B.** 10 nM Cfz. **C.** 15 nM Cfz (AMO-1), or **D.** 10 µM melphalan, compared to DMSO. Red circles are proteins with significantly increased abundances ($p < 0.05$, ≥ 2-fold increase), blue circles are significantly decreased proteins ($p < 0.05$, ≥ 2-fold decrease), and orange circles belong to SRSF family of proteins. Size of dots correspond to summed SILAC light and heavy intensities for each protein. **E.** Volcano plot of $\log_2$ transformed ratio of changes in gene expression for MM.1S treated with 18 nM Cfz, compared to DMSO, with significantly changed genes in green ($p < 0.05$, ≥ 2-fold). **F.** Normalized SILAC LC-MS/MS intensity ratios for phosphopeptides enriched from MM.1S cells treated with 50 µM KH-CB19 versus DMSO for 24 hr. Red and blue circles are significantly changed sites ($p < 0.05$, ≥2-fold). Notably, detected SRSF phosphopeptides (orange circles) do not change significantly in response to CLK1/4 inhibitor. Dot size corresponds to summed SILAC intensities of the phosphopeptides.

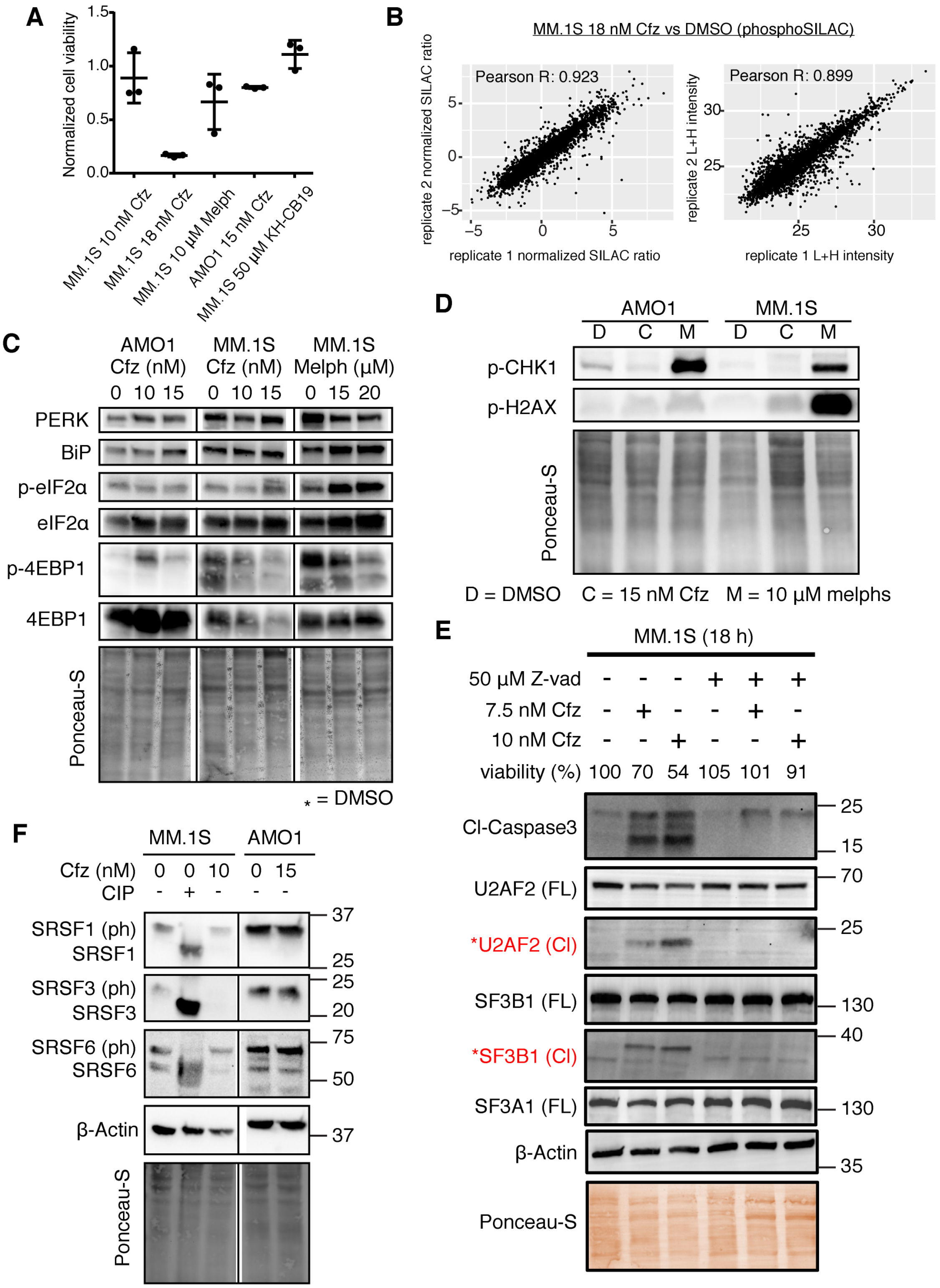

**Figure S3. Characterization of myeloma response to Cfz and melphalan. A.** Normalized cell viability (by CellTiter-Glo) vs. DMSO control of MM cells treated with indicated drugs for 24 hr. (n = 3; mean +/- S.D.). **B.** Representative scatter plot showing correlation between 2 biological replicates for normalized SILAC ratios (left) and summed (light + heavy) intensities (right) for phosphosites from MM.1S treated with 18 nM Cfz shows high quantitative reproducibility. **C.** Immunoblot of stress biomarkers (PERK, BiP, phospho-/total eIF2α, phospho-/total 4EBP1) for AMO-1 and MM.1S treated with DMSO, 10 nM, 15 nM Cfz, and MM.1S treated with DMSO, 15 μM melphalan, and 20 μM melphalan. Vertical lines indicate excised lanes containing conditions not relevant to this study. **D.** Immunoblot of biomarkers for DNA damage (phospho-CHK1, phospho-H2AX) for AMO-1 and MM.1S treated with DMSO, 15 nM Cfz, and 10 μM melphalan, where activation of DNA damage response occurred only with melphalan. **E.** Immunoblot of core spliceosome components, U2AF2, SF3B1, and SF3A1 showing Caspase-3 cleavage (* in red) from MM.1S treated with Cfz for 18 hr with and without 50 μM of Caspase inhibitor zVAD-fmk. Cell viability for each condition after 18 hr is included. **F.** Immunoblot of SRSF1, SRSF3, and SRSF6 in cytoplasmic fraction of MM.1S and AMO-1 treated with DMSO and Cfz. Cell extract treated with calf intestinal phosphatase (+ CIP) to highlight mobility shift of phosphorylated species (ph) of SRSF proteins. Vertical line between cell lines indicate excised lanes containing conditions not relevant to this study. Ponceau-S stain included as loading control for all immunoblots.

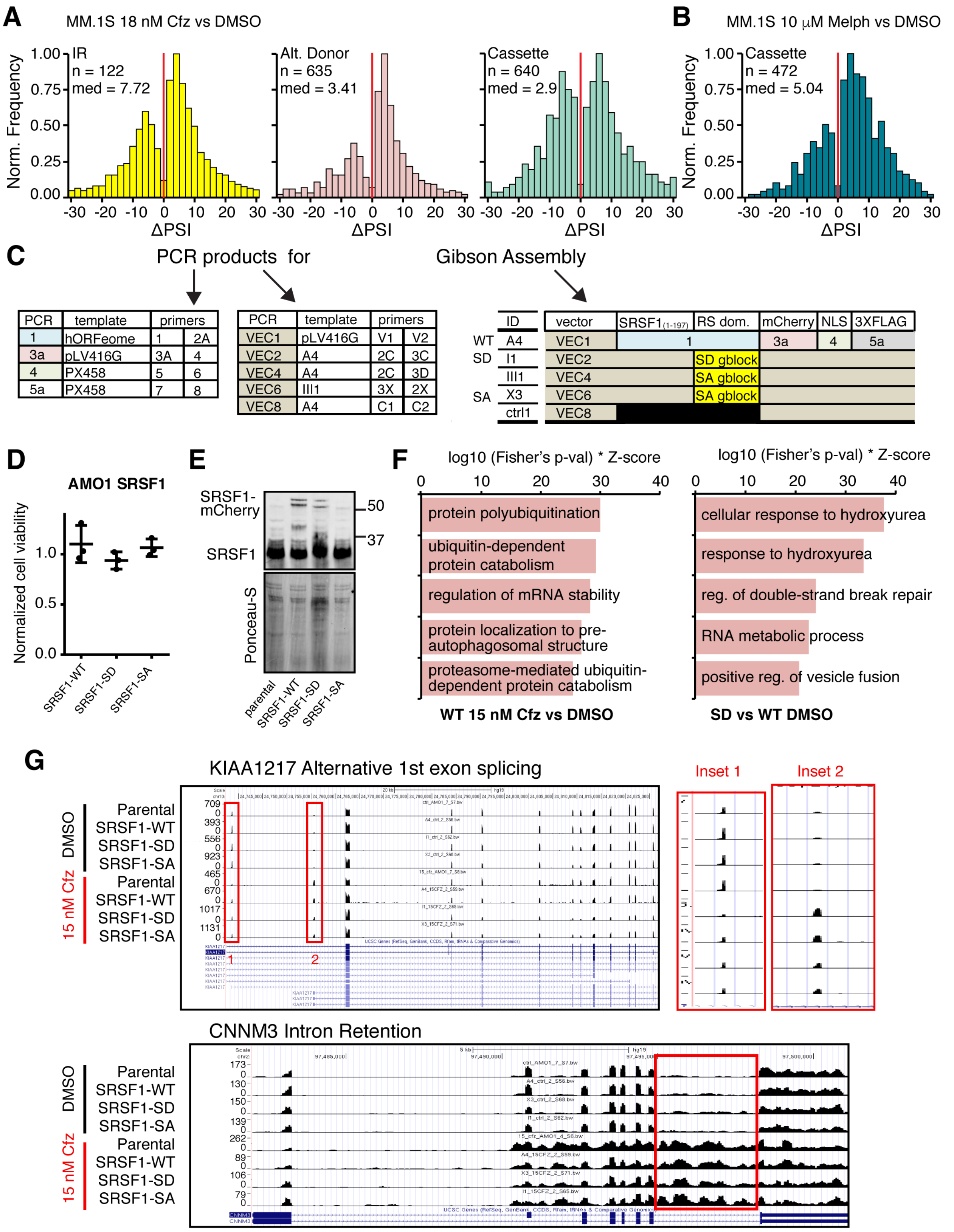
A
MM.1S 18 nM Cfz vs DMSO
B
MM.1S 10 μM Melph vs DMSO
IR
n = 122
med = 7.72
Alt. Donor
n = 635
med = 3.41
Cassette
n = 640
med = 2.9
Cassette
n = 472
med = 5.04
Norm. Frequency
ΔPSI
C
PCR products for
Gibson Assembly
PCR | template | primers
1 | hORFeome | 1 | 2A
3a | pLV416G | 3A | 4
4 | PX458 | 5 | 6
5a | PX458 | 7 | 8
PCR | template | primers
VEC1 | pLV416G | V1 | V2
VEC2 | A4 | 2C | 3C
VEC4 | A4 | 2C | 3D
VEC6 | III1 | 3X | 2X
VEC8 | A4 | C1 | C2
ID | vector | SRSF1(1-197) | RS dom. | mCherry | NLS | 3XFLAG
WT A4 VEC1 1 3a 4 5a
SD I1 VEC2 SD gblock
III1 VEC4 SA gblock
SA X3 VEC6 SA gblock
ctrl1 VEC8
D
AMO1 SRSF1
Normalized cell viability
SRSF1-WT
SRSF1-SD
SRSF1-SA
E
SRSF1-mCherry
SRSF1
50
37
Ponceau-S
parental
SRSF1-WT
SRSF1-SD
SRSF1-SA
F
log10 (Fisher's p-val) * Z-score
protein polyubiquitination
ubiquitin-dependent protein catabolism
regulation of mRNA stability
protein localization to pre-autophagosomal structure
proteasome-mediated ubiquitin-dependent protein catabolism
WT 15 nM Cfz vs DMSO
log10 (Fisher's p-val) * Z-score
cellular response to hydroxyurea
response to hydroxyurea
reg. of double-strand break repair
RNA metabolic process
positive reg. of vesicle fusion
SD vs WT DMSO
G
KIAA1217 Alternative 1st exon splicing
Inset 1
Inset 2
DMSO
15 nM Cfz
Parental
SRSF1-WT
SRSF1-SD
SRSF1-SA
CNNM3 Intron Retention

**Figure S4. Cfz-induced splicing alterations across parental MM cells and SRSF constructs. A.** and **B.** ΔPSI histograms of significant ($p < 0.05$) IR, alt. exon donor, and alt. cassette splicing events for MM.1S treated with 18 nM Cfz for 24 hr (or **B.** 10 µM melphalan) with respect to DMSO. Subset of data in **Fig. 3B, D**. Bin = 2 and red line at ΔPSI = 0. **C.** List of PCR reactions referencing oligos in **Supplementary Table S1**.and the subsequent Gibson Assemblies to generate lentiviral SRSF1 plasmids. IDs are related to SRSF1-WT and mutants: A4 = WT, I1 = SD, III1/X3 = SA. **D.** Normalized cell viability of AMO-1 cells expressing SRSF1-WT, SRSF1-SD, and SRSF1-SA treated with 15 nM Cfz for 24 hr ($n = 3$; mean +/- S.D.). **E.** SRSF1 immunoblot of parental AMO-1, AMO-1 expressing SRSF1-WT, SRSF1-SD, and SRSF1-SA, comparing exogenous SRSF1-mCherry-NLS-$[FLAG]_3$ and endogenous SRSF1 abundance. **F.** Top ranked (combined Fisher's p-value and background weighted Z-score) GO enrichment terms for genes with significant ASE ($p < 0.05$) of all types from WT treated with 15 nM Cfz compared to DMSO (left) and SD compared to WT in DMSO (right). **G.** Examples of Cfz-induced alternative splicing: alternative first exon splicing in *KIAA 1217* (top) and intron retention in *CNNM3* (bottom) for all AMO-1 cells (parental, SRSF1-WT, SD, SA) with 15 nM Cfz compared to DMSO.

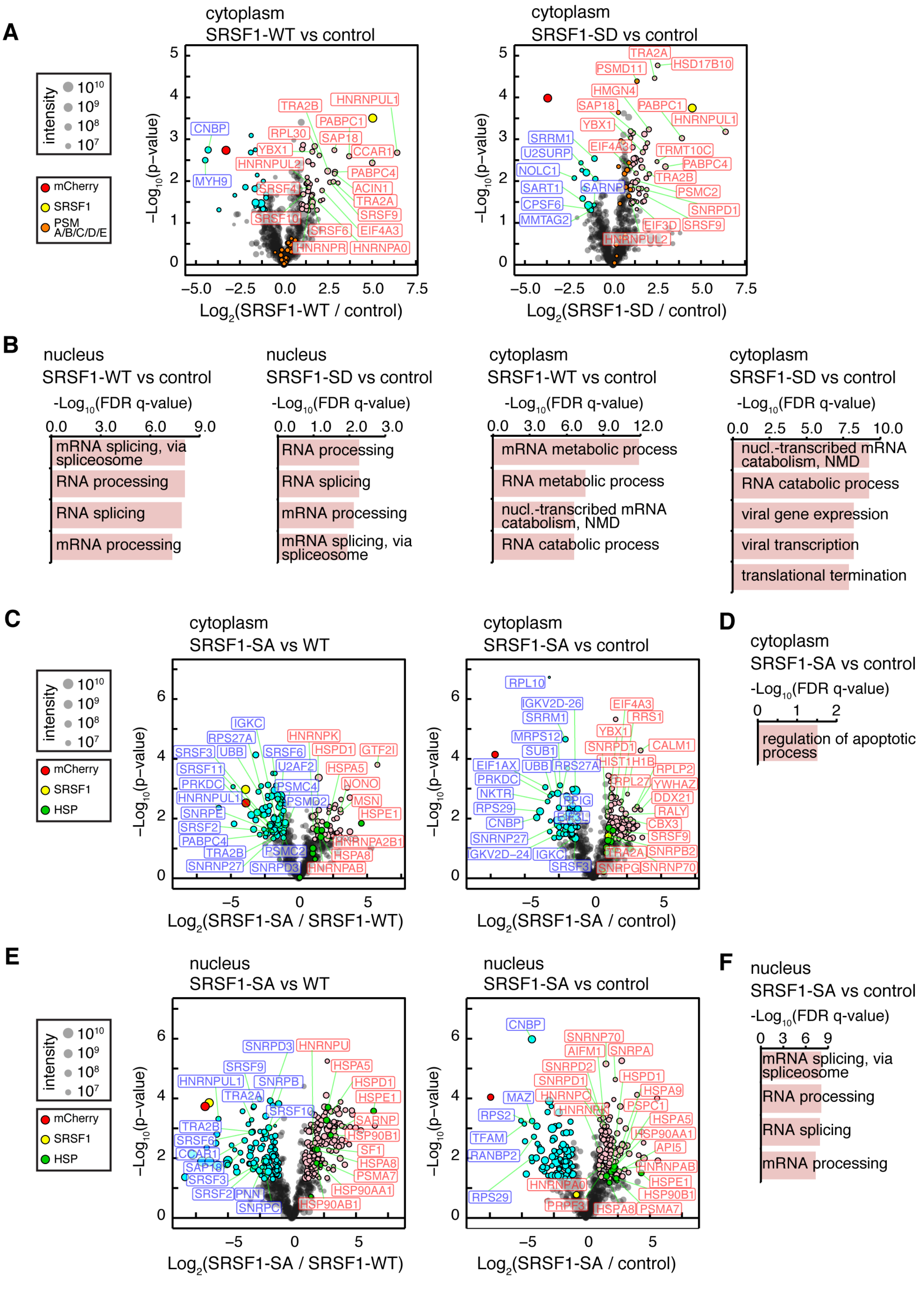
A
cytoplasm
SRSF1-WT vs control
cytoplasm
SRSF1-SD vs control
intensity
mCherry
SRSF1
PSM A/B/C/D/E
−Log10(p−value)
Log2(SRSF1-WT / control)
Log2(SRSF1-SD / control)
B
nucleus
SRSF1-WT vs control
-Log10(FDR q-value)
mRNA splicing, via spliceosome
RNA processing
RNA splicing
mRNA processing
nucleus
SRSF1-SD vs control
RNA processing
RNA splicing
mRNA processing
mRNA splicing, via spliceosome
cytoplasm
SRSF1-WT vs control
mRNA metabolic process
RNA metabolic process
nucl.-transcribed mRNA catabolism, NMD
RNA catabolic process
cytoplasm
SRSF1-SD vs control
nucl.-transcribed mRNA catabolism, NMD
RNA catabolic process
viral gene expression
viral transcription
translational termination
C
cytoplasm
SRSF1-SA vs WT
cytoplasm
SRSF1-SA vs control
HSP
Log2(SRSF1-SA / SRSF1-WT)
Log2(SRSF1-SA / control)
D
cytoplasm
SRSF1-SA vs control
regulation of apoptotic process
E
nucleus
SRSF1-SA vs WT
nucleus
SRSF1-SA vs control
F
nucleus
SRSF1-SA vs control
mRNA splicing, via spliceosome
RNA processing
RNA splicing
mRNA processing

**Figure S5. The phosphorylation-dependent interactome of SRSF1. A.** Volcano plot showing AP-MS enriched interaction partners to SRSF1-WT (*left*) and SRSF1-SD (*right*) compared with mCherry-NLS-$[FLAG]_3$ control in AMO-1 cytoplasm. Significantly enriched proteins ($p < 0.05$, ≥ 2-fold) are in pink and excluded proteins in cyan. Dot size corresponds to combined LFQ intensities for that protein. Proteasomal subunits are colored in orange. **B.** Top ranked (FDR *q*-value) GO enrichment terms for significantly enriched interaction partners for SRSF1-WT and SRSF1-SD compared to mCherry-NLS-$[FLAG]_3$ control in either the nucleus or cytoplasm of AMO-1. **C and E**. Volcano plots depicting differential interaction partners of SRSF1-SA compared to SRSF1-WT (*left*) and SRSF1-SA enriched proteins compared to control (*right*) in **C)** the cytoplasm and **E)** the nucleus of AMO-1. Significantly enriched and excluded proteins ($p < 0.05$, ≥ 2-fold) in pink and cyan. Heat shock proteins colored in green. **D and F.** Top ranked (FDR *q*-value) GO enrichment terms for significantly enriched interaction partners for SRSF1-SA in either the **D)** cytoplasm or **F)** nucleus of AMO-1.

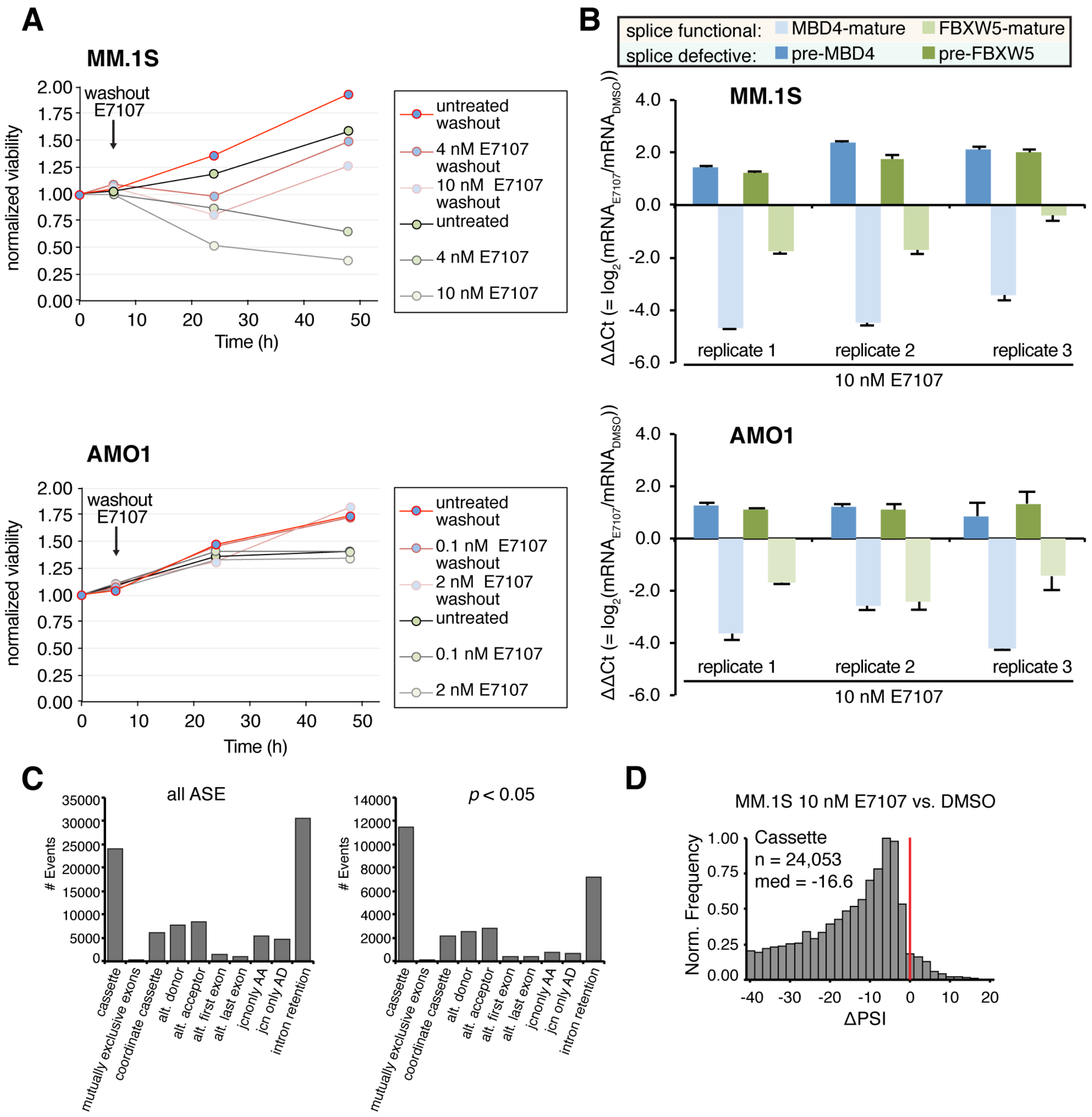

**Figure S6. E7107 cell toxicity, functional splicing assay, and splicing statistics. A.** Normalized cell viability timecourse for MM.1S (top) and AMO-1 (bottom) treated with increasing E7107 concentration. Cells with a washout of E7107 (with exchanged media) at 6 hr have pink and red outlines, while cells without washout are shown with gray and black outlines. **B.** Bar graph depicting $\log_2$ transformed ratio of transcript abundance between cells treated with 10 nM E7107 for 6 hr, and DMSO, via qPCR ΔΔCt values (normalized to housekeeping gene *PPIA*) of pre-splice and splice competent forms of two targets (*MBD4* in blue and *FBXW5* in green) in MM.1S (top) and AMO-1 cells confirm significant intron retention of canonical targets at this dose and time point. Error bars represent standard deviation of technical replicates ($n$ = 3). **C.** Distribution of all JuncBASE quantified splice events (n = 89,988) across the ASE types (left) and only significant ($p < 0.05$) events (n = 28,436) (right) in MM.1S treated with 10 nM E7107 for 6 hr compared to DMSO shows IR and alternative cassette exon splicing events are the most common types, regardless of significance. **D.** ΔPSI histogram of cassette events for MM.1S treated with 10 nM E7107 for 6 hr with respect to DMSO shows the effect of impaired splicing on exon selection. Bin = 2 and red line at ΔPSI = 0.

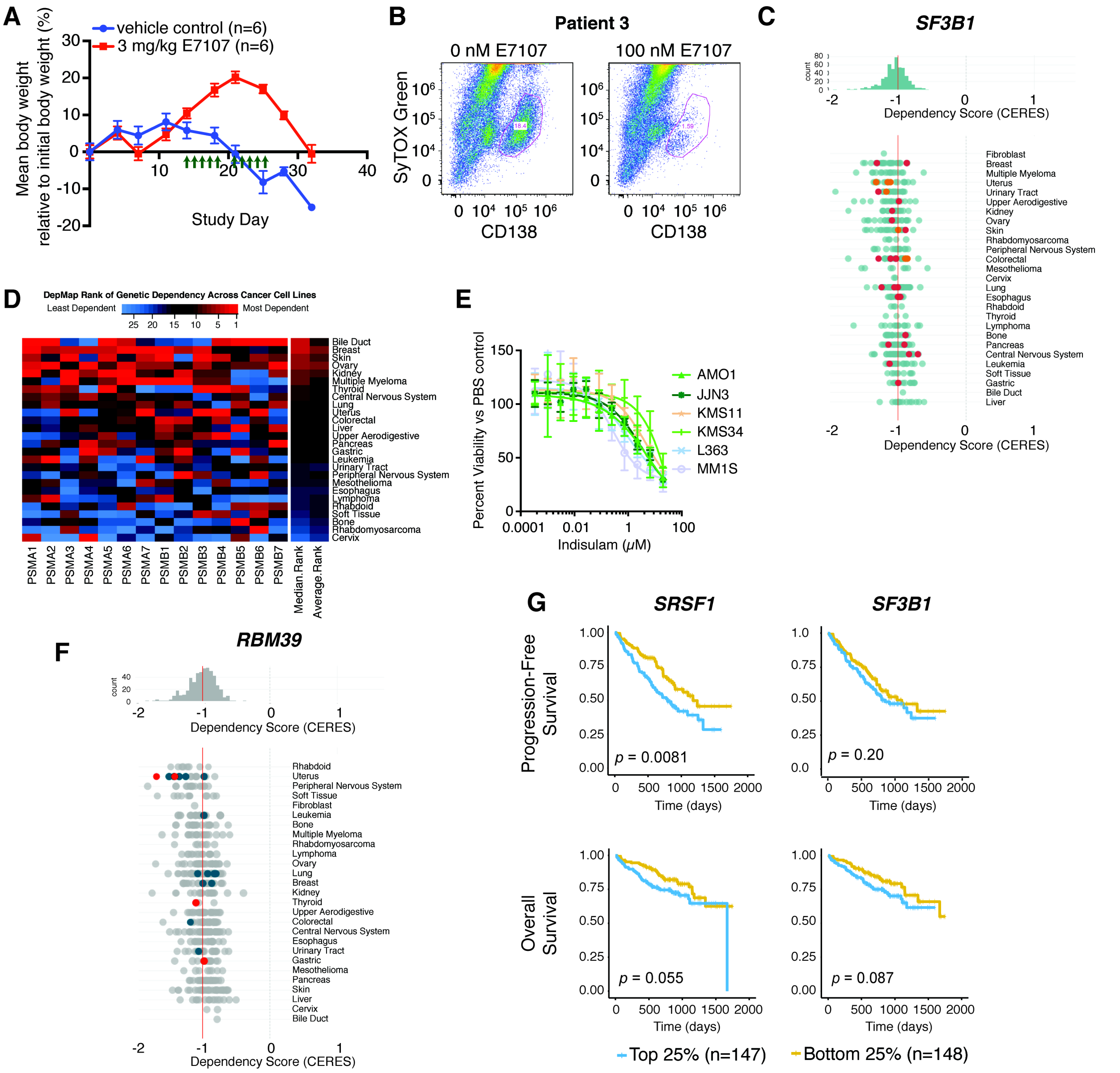

**Figure S7. Preclinical and clinical relevance of targeting the spliceosome in myeloma. A.** 3 mg/kg I.V. E7107 leads to minimal weight loss in NSG mice (mean +/- S.D.; *n* = 6 per arm). **B.** Example flow cytometry dot plots from primary MM patient bone marrow aspirates. Gated region indicates CD138+ plasma cells. **C.** DepMap (www.depmap.org) CRISPR screen (**Avana** library **18Q4**) dependency data indicating that MM cell lines are among the most sensitive to *SF3B1* depletion based on cell line rank (1 = most sensitive, with lowest average CERES score; 27 = least sensitive, with highest average CERES score). Red line indicates cutoff (CERES score = -1) indicative of an essential gene. **D.** Aggregated DepMap ranking of cancer cell lines across all genes comprising the 20S proteasome, indicating increased but not maximal sensitivity of MM lines to genetic proteasome subunit ablation. **E.** Cytotoxicity of indisulam versus a panel of MM cell lines (48 hr treatment; $n = 4$ per data point, +/- S.D.). **F.** DepMap dependency data (as in **D.**) for RBM39. **G.** Progression-free and overall survival data from the CoMMpass study (version **IA11**) with respect to top and bottom quartiles of *SRSF1* and *SF3B1* gene expression in newly-diagnosed MM patient tumor cells. *p*-values by log-ranked test.

**Table S1: Oligo sequences for cloning, qPCR, and RNA-seq cDNA library.** Oligo ID for CLONING

| | oligo ID | sequence | target gene | vendor |
|---|---|---|---|---|
| CLONING | V1 | AAGATGGCCTAGTAGACCCAGCTTTCTTGTACAAAGTGG | pLV416G | sigma |
| | V2 | CCCGACATAGCAGCCTGCTTTTTTGTACAA | pLV416G | sigma |
| | 1 | GGCTGCTATGTCGGGAGGTGGTGTGAT | SRSF1 | sigma |
| | 2A | CTGGTGCCTGTACGAGAGCGAGATCTGCTATGAC | SRSF1 | sigma |
| | 3A | TCGTACAGGCACCAGCGGTACCA | mCherry | sigma |
| | 4 | CGGCCTTTTCTTGTACAGCTCGTCCATGCC | mcherry | sigma |
| | 5 | TGTACAAGAAAAGGCCAGCGGCCAC | NLS | sigma |
| | 6 | CCTTATAGTCGCCGAATTCCTTTTTCTTTTTTGCCT | NLS | sigma |
| | 7 | GAATTCGGCGACTATAAGGACCACGACGGAGACT | 3XFLAG | sigma |
| | 8 | GGTCTACTAGGCCATCTTATCGTCATCGTCTTTG | 3XFLAG | sigma |
| | 2C | CATCAACTTTAACCCGGATGTAGGC | SRSF1 | sigma |
| | 3C | TCGTACAGGCACCAGCGGTACCA | mCherry | sigma |
| | 3D | CCCGTACAATGGTGAGCAAGGGCGAGG | mCherry | sigma |
| | 4B | CCTTATAGTCCTTGTACAGCTCGTCCATGCC | mCherry | sigma |
| | 7B | GCTGTACAAGGACTATAAGGACCACGACGGAG | 3XFLAG | sigma |
| | 3X | GGCACCAGCGGTACAATGGTGAGCAAGGGCGAGGAGG | mCherry | sigma |
| | 2X | TGTACCGCTGGTGCCTGTACGGGCGCGGGCTCTG | SRSF1 | sigma |
| | C1 | AGGCTGCTATGGTGAGCAAGGGCGAG | pLV416G | sigma |
| | C2 | CTCACCATAGCAGCCTGCTTTTTTGTACAAACT | pLV416G | sigma |
| | SD gblock | ATCCGGGTTAAAGTTGATGGGCCCAGAGATCCAGATTATGGAAGAGATCGAGATCGAGATCGTGATCGTGATAGAGATCGTGATAGAGATAA CGATAGGGATCGCGATTACGATCCAAGGAGAGATAGAGGAGATCCACGCTATGATCCCCGTCATGATAGAGATCGCGATCGTACAGGCACCAG | | IDT |
| | SA gblock | ATCCGGGTTAAAGTTGATGGGCCCAGAGCCCCAGCCTATGGAAGAGCCCGAGCCCGAGCCCGTGCCCGTGCCAGAGCCCGTGCCAGAGCCAAC GCCAGGGCCCGCGCCTACGCTCCAAGGAGAGCCAGAGGAGCTCCACGCTATGCTCCCCGTCATGCCAGAGCCCGCGCCCGTACAATGGTGAG | | IDT |
| qPCR | F1 | CACACCAGATCGGCATCAA | FBXW5 mature | sigma |
| | F2 | CGATGATGTGTCCGTGTATGT | FBXW5 mature | sigma |
| | F3 | AGACACCACTGAGGTAGGAA | FBXW5 unspliced | sigma |
| | F4 | TGGCAGGATCTGCTTGATG | FBXW5 unspliced | sigma |
| | M1 | GGAAGCTTCTCATCGCTACTAT | MBD4 | sigma |
| | M2 | GTTCTGACACATCTCTCCAGTC | MBD4 mature | sigma |
| | M3 | CCCTACCACACTGTCTCTACTA | MBD4 unspliced | sigma |
| | PP1 | AGACAAGGTCCCAAAGAC | PPIA | sigma |
| | PP2 | ACCACCCTGACACATAAA | PPIA | sigma |
| TruSeq adapters | Univ. Adapt. (P5) | AATGATACGGCGACCACCGAGATCTACACTCTTTCCCTACACGACGCTCTTCCGATC*T | [* = phosphorothioate] | sigma |

| | | | | |
|---|---|---|---|---|
| P7 (barcode sequence under-lined) | Index 1 | GATCGGAAGAGCACACGTCTGAACTCCAGTCACATCACGATCTCGTATGCCGTCTTCTGCTTG | | sigma |
| | Index 2 | GATCGGAAGAGCACACGTCTGAACTCCAGTCACCGATGTATCTCGTATGCCGTCTTCTGCTTG | | sigma |
| | Index 3 | GATCGGAAGAGCACACGTCTGAACTCCAGTCACTTAGGCATCTCGTATGCCGTCTTCTGCTTG | | sigma |
| | Index 4 | GATCGGAAGAGCACACGTCTGAACTCCAGTCACTGACCAATCTCGTATGCCGTCTTCTGCTTG | | sigma |
| | Index 5 | GATCGGAAGAGCACACGTCTGAACTCCAGTCACACAGTGATCTCGTATGCCGTCTTCTGCTTG | | sigma |
| | Index 6 | GATCGGAAGAGCACACGTCTGAACTCCAGTCACGCCAATATCTCGTATGCCGTCTTCTGCTTG | | sigma |
| | Index 7 | GATCGGAAGAGCACACGTCTGAACTCCAGTCACCAGATCATCTCGTATGCCGTCTTCTGCTTG | | sigma |
| | Index 8 | GATCGGAAGAGCACACGTCTGAACTCCAGTCACACTTGAATCTCGTATGCCGTCTTCTGCTTG | | sigma |
| | Index 9 | GATCGGAAGAGCACACGTCTGAACTCCAGTCACGATCAGATCTCGTATGCCGTCTTCTGCTTG | | sigma |
| | Index 10 | GATCGGAAGAGCACACGTCTGAACTCCAGTCACTAGCTTATCTCGTATGCCGTCTTCTGCTTG | | sigma |
| | Index 11 | GATCGGAAGAGCACACGTCTGAACTCCAGTCACGGCTACATCTCGTATGCCGTCTTCTGCTTG | | sigma |
| | Index 12 | GATCGGAAGAGCACACGTCTGAACTCCAGTCACCTTGTAATCTCGTATGCCGTCTTCTGCTTG | | sigma |
| | Index 13 | GATCGGAAGAGCACACGTCTGAACTCCAGTCACAGTCAAATCTCGTATGCCGTCTTCTGCTTG | | sigma |
| | Index 14 | GATCGGAAGAGCACACGTCTGAACTCCAGTCACAGTTCCATCTCGTATGCCGTCTTCTGCTTG | | sigma |
| | Index 15 | GATCGGAAGAGCACACGTCTGAACTCCAGTCACATGTCAATCTCGTATGCCGTCTTCTGCTTG | | sigma |
| | Index 16 | GATCGGAAGAGCACACGTCTGAACTCCAGTCACCCGTCCATCTCGTATGCCGTCTTCTGCTTG | | sigma |
| | Index 18 | GATCGGAAGAGCACACGTCTGAACTCCAGTCACGTCCGCATCTCGTATGCCGTCTTCTGCTTG | | sigma |
| | Index 19 | GATCGGAAGAGCACACGTCTGAACTCCAGTCACGTGAAAATCTCGTATGCCGTCTTCTGCTTG | | sigma |

**Table S2. Genetic constructs.** Description of SRSF1 constructs

| ID | Name | Features | Promoter | Details |
|---|---|---|---|---|
| **A4** | SRSF1 WT | SRSF1-mCherry-NLS-3XFLAG | EF1α | human SRSF1 was obtained from hORFeome v8.1; plasmid backbone was amplified by PCR from pLV416G-f-luc/mCherry lentiviral vector used for constitutive expression of luciferase in mouse models |
| **I1** | SRSF1mSD | SRSF1(1-197)-all RS domain S--> D substitution-mCherry-NLS-3XFLAG | EF1α | templated from A4 construct |
| **X3** | SRSF1mSA | SRSF1(1-197)-all RS domain S--> A substitution-mCherry-NLS-3XFLAG | EF1α | templated from III1 construct, which was initial SRSF1mSA construct with shortened mCherry linker |
| **ctrl1** | ctrl1 | mCherry-NLS-3XFLAG | EF1α | templated from A4 construct |

**Table S3. JuncBASE statistics.** Distribution of ΔPSI calculated in R. Statistics for individual event types and all event types are listed for each comparative condition

| Condition | event_type | total events | significant events (raw_pval <0.05) | significant events (raw_pval < 0.05) & \|deltaPSI\| ≥ 10 | median deltaPSI total | median deltaPSI significant (raw_pval < 0.05) | Mode total (bin size = 2) | Mode sig. (raw_pval < 0.05, bin size = 2) |
|---|---|---|---|---|---|---|---|---|
| MM.1S 18 nM Cfz vs. MM.1S DMSO | cassette | 11125 | 640 | 254 | 0.29 | 2.895 | 2 | 6 |
| | mutually_exclusive | 68 | 2 | 1 | -2.355 | -12.915 | -6 | -20;6 |
| | coord_cassette | 1164 | 43 | 23 | 0.54 | 5.56 | 2 | 16 |
| | alternative_donor | 9188 | 635 | 215 | 1.64 | 3.41 | 2 | 4 |
| | alternative_acceptor | 9352 | 630 | 167 | 0.81 | 3.035 | 2 | 2 |
| | alternative_first_exon | 1443 | 100 | 50 | 0.3 | 2.71 | 0 | 2; 6 |
| | alternative_last_exon | 1145 | 73 | 32 | -0.1 | 1.9 | 0 | 6 |
| | jcn_only_AA | 6759 | 284 | 103 | 0 | 1.645 | 2 | 4 |
| | jcn_only_AD | 5595 | 277 | 129 | 0.09 | -1.31 | 2 | -8; 10 |
| | intron_retention | 25807 | 122 | 69 | 2.54 | 7.72 | 2 | 6 |
| | **All Types** | **71646** | **2806** | **1043** | **1.52** | **3.07** | **2** | **4** |
| MM.1S 10 uM melph vs. MM.1S DMSO | cassette | 12267 | 472 | 187 | 1.06 | 5.035 | 2 | 4 |
| | mutually_exclusive | 84 | 2 | 1 | -1.455 | -8.39 | -2 | -22; 6 |
| | coord_cassette | 1417 | 53 | 23 | 1.75 | 6.52 | 2 | 4; 8 |
| | alternative_donor | 9535 | 331 | 91 | 0.37 | 2.33 | 0 | 2 |
| | alternative_acceptor | 10445 | 280 | 68 | 0.18 | 1.965 | 0 | 4 |
| | alternative_first_exon | 1561 | 85 | 37 | 0.37 | 2.41 | 0 | 6 |
| | alternative_last_exon | 1328 | 58 | 18 | -0.215 | 0.685 | 0 | -4 |
| | jcn_only_AA | 7667 | 240 | 87 | -0.13 | -2.48 | -2 | -4 |
| | jcn_only_AD | 6651 | 294 | 136 | -0.14 | -1.91 | -2 | -8 |
| | intron_retention | 24247 | 24 | 16 | 0.44 | -6.07 | 0 | -32; -28 |
| | **All Types** | **75202** | **1839** | **664** | **0.42** | **2.87** | **0** | **4** |
| AMO-1 15 | cassette | 11645 | 685 | 313 | -0.82 | -4.72 | -2 | -4 |

| | | | | | | | |
|---|---|---|---|---|---|---|---|
| nM Cfz vs. AMO-1 DMSO | mutually_exclusive | 397 | 15 | 5 | -0.65 | -7.48 | -6 | -8; 6 |
| | coord_cassette | 1673 | 58 | 33 | -1.37 | -5.41 | -4 | 6 |
| | alternative_donor | 8116 | 358 | 147 | 0.605 | 3.83 | 2 | 4 |
| | alternative_acceptor | 9620 | 385 | 128 | 0.4 | 3.7 | 2 | 4 |
| | alternative_first_exon | 1241 | 90 | 53 | 0.13 | 2.465 | -2; 0 | 8 |
| | alternative_last_exon | 899 | 36 | 21 | 0.05 | 5.075 | 0 | 12 |
| | jcn_only_AA | 10700 | 368 | 165 | 0.2 | 3.185 | 2 | 6 |
| | jcn_only_AD | 9795 | 390 | 196 | 0.3 | -2.125 | 2 | 6 |
| | intron_retention | 27386 | 174 | 126 | 2.2 | 12.835 | 2 | 6 |
| | **All Types** | **81472** | **2559** | **1187** | **0.77** | **2.9** | **2** | **6** |
| MM.1S 10 nM E7107 vs. MM.1S DMSO | cassette | 24053 | 11484 | 9945 | -16.6 | -31.66 | -6 | -8 |
| | mutually_exclusive | 91 | 21 | 16 | -0.5 | 5.64 | 0 | 6; 10 |
| | coord_cassette | 6137 | 2200 | 1904 | -13.34 | -34.09 | -4 | -6 |
| | alternative_donor | 7854 | 2502 | 1618 | 3.43 | 8.545 | 4 | 4 |
| | alternative_acceptor | 8493 | 2835 | 1846 | 3.09 | 7.59 | 4 | 6 |
| | alternative_first_exon | 1402 | 377 | 267 | -1.38 | -5.45 | -2 | -12 |
| | alternative_last_exon | 1088 | 356 | 245 | -3.705 | -9.8 | -2 | -8; -6 |
| | jcn_only_AA | 5495 | 792 | 463 | -0.02 | -2.24 | 2 | -6 |
| | jcn_only_AD | 4709 | 698 | 415 | 0.02 | 2.805 | 4 | 8 |
| | intron_retention | 30666 | 7171 | 6564 | 13.79 | 37.2 | 6 | 24 |
| | **All Types** | **89988** | **28436** | **23283** | **1.27** | **-8.685** | **4** | **-6** |
| SRSF1 WT 15 nM Cfz vs. SRSF1 WT DMSO | cassette | 8337 | 539 | 215 | 0.48 | 2.56 | 2 | 10 |
| | mutually_exclusive | 147 | 3 | 1 | -0.42 | -2.9 | 2 | -20; -2 |
| | coord_cassette | 798 | 51 | 25 | 1.38 | 6.7 | 4 | 10 |
| | alternative_donor | 7639 | 343 | 98 | 0.83 | 3.74 | 2 | 10 |
| | alternative_acceptor | 8410 | 450 | 104 | 0.64 | 3.255 | 0 | 5 |
| | alternative_first_exon | 1285 | 112 | 63 | 0.55 | 5.11 | 0 | 10 |
| | alternative_last_exon | 1183 | 80 | 32 | 0.41 | 4.17 | 0 | 10 |
| | jcn_only_AA | 7030 | 318 | 112 | 0.035 | -0.845 | -2 | -25 |
| | jcn_only_AD | 6257 | 371 | 160 | 0.32 | 3.06 | 2 | 10 |

| | | | | | | | | |
|---|---|---|---|---|---|---|---|---|
| | intron_retention | 12139 | 60 | 36 | 2.45 | 7.865 | 2 | 10 |
| | **All Types** | **53225** | **2327** | **846** | **1.09** | **3.24** | **2** | **4** |
| SRSF1mSD DMSO vs. SRSF1 WT DMSO | cassette | 8425 | 335 | 94 | 0.09 | 1.79 | 0 | 4 |
| | mutually_exclusive | 184 | 4 | 0 | 0.025 | 1.025 | 4 | -2 |
| | coord_cassette | 810 | 39 | 22 | 0.12 | 2.83 | -2 | 4 |
| | alternative_donor | 7204 | 217 | 45 | 0.05 | -0.28 | 0 | 2 |
| | alternative_acceptor | 8192 | 209 | 43 | 0.07 | 1.37 | 0 | 4 |
| | alternative_first_exon | 1345 | 60 | 20 | 0.02 | 0.995 | 0 | -6 |
| | alternative_last_exon | 1175 | 38 | 11 | 0.21 | 2.98 | 0 | 6 |
| | jcn_only_AA | 7139 | 234 | 100 | 0 | 1.66 | 0 | -4 |
| | jcn_only_AD | 6247 | 237 | 98 | 0 | 1.95 | 2 | 4 |
| | intron_retention | 10643 | 33 | 19 | 0.17 | -12.5 | 0 | -10 |
| | **All Types** | **51364** | **1406** | **452** | **0.08** | **1.1** | **0** | **4** |
| **Conditions** | event_type | **total events** | **significant events (corrected_pval <0.05)** | **significant events (corrected_pval < 0.05) & \|deltaPSI\| ≥ 10** | **median deltaPSI total** | **median deltaPSI significant (corrected_pval < 0.05)** | **Mode_total (bin size = 2)** | **Mode_significant (corrected_pval < 0.05, bin size = 2)** |
| AMO-1 (all: parental, WT, SD, SA) 15 nM Cfz vs. AMO-1 (all) DMSO | cassette | 9401 | 286 | 115 | 0.08 | -3.49 | 0 | 5 |
| | mutually_exclusive | 47 | 0 | 0 | 0.42 | NA | 0 | 0 |
| | coord_cassette | 562 | 6 | 3 | 0.425 | 8.775 | 0 | 5 |
| | alternative_donor | 8147 | 1134 | 139 | 0.68 | 1.83 | 0 | 10 |
| | alternative_acceptor | 9000 | 810 | 114 | 0.54 | 1.635 | 0 | 10 |
| | alternative_first_exon | 1260 | 119 | 58 | 0.28 | 3.69 | 0 | 10 |
| | alternative_last_exon | 1217 | 61 | 21 | 0 | 1.24 | 0 | 5 |
| | jcn_only_AA | 5482 | 48 | 21 | 0 | 1.705 | 0 | -5 |
| | jcn_only_AD | 4799 | 68 | 24 | 0.07 | -1.95 | 0 | 5 |
| | intron_retention | 22559 | 43 | 25 | 1.53 | 10.8 | 0 | 10 |
| | **All Types** | **62474** | **2575** | **520** | **0.84** | **1.76** | **0** | **2** |

### Supplementary Methods

#### Cell lines

MM.1S is from ATCC (CRL-2974), AMO-1, AMO1 Btz-resistant are gifts courtesy of Dr. Christoph Driesen (Kantonsspital St Gallen), L363, RPMI8266, JJN3 are from Deutsche Sammlung von Mikroorganismen und Zellkulturenrepository (DSMZ; ACC 49, ACC 402, ACC 541 respectively), INA6 is a gift courtesy of Dr. Renate Burger (Christian-Albrechts-Universität zu Kiel), KMS34 is from Japanese Collection of Research Bioresources Cell Bank (JCRB1195), and MM.1S mCherry/f-luc was provided by UCSF Hematologic Malignancies Tissue Bank (HMTB). All cell lines used are female, except INA6 and RPMI8266. All cell lines were validated using STR profiling service by ATCC.

#### Cell culture for PI-response proteomics

For single-timepoint experiments, cells were passaged in either light SILAC media (SILAC RPMI 1640 (Thermo, PI88421), supplemented with 1% Pencillin/Streptomycin, 10% dialyzed FBS (Thermo, PI88440), and 321.6 µM L-Lysine and 190.4 µM L-Arginine (Sigma, L8662-25G, A6969-25G) or heavy SILAC media (L-Lysine-$^{13}C_6$,$^{15}N_2$, L-Arginine-$^{13}C_6$,$^{15}N_4$ (Cambridge Isotope, CNLM-291-H-1, CNLM-539-H-1) instead of L-Lys and L-Arg) for more than 6 doublings previous to drug dosing experiments to mostly incorporate heavy and light lysine and arginine. For SRSF1 AP-MS, ~30E+6 cells were treated in complete media with 15 nM Cfz or DMSO for 24 hr. Cells were harvested by centrifugation at 300 rcf for 5 min and washed with 5 mL PBS. Cells were then pelleted by centrifugation, PBS was aspirated, and cell pellets were frozen in liquid nitrogen (LN2) and stored in -80°C.

#### E7107 timecourse and washout

For E7107 time-course “washout” experiment in **Fig. S6A**, cells are treated in 6-well plates at 1E+6 cells/mL with DMSO, 0.1 nM, 2 nM E7107 for AMO-1 and DMSO, 4 nM , and 10 nM E7107 for MM.1S . Cell viability was measured with CellTiter-Glo (Promega, G7573) at 0 hr, 6 hr, 24 hr, and 48 hr, where media for “washout” samples were exchanged for fresh media, without E7107after the 6 hr measurement.

**SRSF1 Cloning details**

**Refer to Supplementary Table S1** for primer sequences and **Table S2 and Fig. S4C** for construct details and assembly of lentiviral vectors encoding SRSF1-mCherry-(NLS)-$[FLAG]_3$ and its RS domain mutants used in SRSF1 AP-MS experiments, microscopy experiments, and AMO-1 exogenous SRSF1 alternative splicing analysis.

Plasmid encoding SRSF1 was isolated from Human ORFeome library v8.1 (1) (Access provided by UCSF Recombinant Antibody Network), and polymerase chain reaction (PCR) was used to amplify the gene from this plasmid, nucleoplasmin nuclear localization signal (NLS) and 3X FLAG sequences from PX458, and pLV lentiviral transfer plasmid backbone and mCherry from pLV-416G. Gibson assembly was used to combine SRSF1, mCherry, NLS, and 3X FLAG with the lentiviral vector backbone (excluding luciferase-T2A-mCherry genes) pLV-416G and this WT construct is referred to as A4. SRSF1 mutant constructs were formed by PCR amplification of a truncated SRSF1 (1-197) in A4, and Gibson assembly with gBlock oligos (IDT), synthetically designed oligos substituting native codons for all Ser with codons for Asp or Ala, depending on it being a SD or SA mutant. SD construct is referred to as I1, while SA construct is referred to as III1. However, SA gBlock design left a shortened linker region. This was corrected by further PCR amplification of III1 plasmid with extended primers to create corrected SA referred to as X3. AP-MS negative control of mCherry-(NLS)-$[FLAG]_3$, referred to as ctrl1, was constructed by PCR amplification of A4 with primers excluding SRSF1 and annealed by one fragment Gibson Assembly. Template vector was removed by DpnI nuclease treatment.

**Lentiviral transduction details**

For each sample, 1.5 µg lentiviral-SRSF1 transfer plasmid was transfected along with 1.33 µg of the packaging plasmid pCMV-dR8.91 (containing Gag-Pol) and 0.17 µg of the VSV-G envelope expressing plasmid pMD2.G into Lenti-X (Takara, 632180) packaging cells (seeded the day before in 6-well plates with 0.6E+6 cells and 2.6 mL Opti-MEM (Life Tech, 31985-062) per well) with FuGene (Promega, E2311) transfection reagent in 300 µl Opti-MEM, incubating for 30min, before adding to each well. After 2 days transfection, viral particles were harvested and filtered with 0.45 µm filter and concentrated with 1 part viral titer and 3 part Lenti-X concentrator (Takara, 631231, ~9-10 mL total) by incubating at 4°C for more than 12 hr, then spinning at 1500 rcf for 45 min at 4°C. Supernatant is carefully aspirated and virus is resuspended in PBS. Entire viral titers were distributed between AMO-1 and MM.1S cells. ~0.75-1.5E+6 cells were seeded in each well of a 6-well plate with 1.5 mL normal growth media, with 8 µg/mL polybrene added (final concentration of 4 µg/mL) and 1mL of virus and 0.5 mL media, then mixed together. Cells were transduced by spinfection, spinning plates at 1000 rcf at 33°C for 2 hr. Afterwards, plates are stored in 37°C incubator (5% $CO_2$) for 2 days, before media is replaced. After a few passages, positively transduced cells with G418 (VWR, 970 3-058), for several passages and then sorted for mCherry fluorescence by Fluorescence Activated Cell Sorting (FACS, Sony SH800). Cells were maintained as all other MM cell lines.

**LC-MS/MS settings**

All samples were analyzed by means of a 3h 15 min non-linear gradient from 2.4% acetonitrile (ACN), 0.1% FA to 32% ACN, 0.1% FA, at 0.2 µL/min, 6 min linear gradient to 79% ACN, 0.1% FA at 0.5 µL/min, then washed with flowrate 0.5 µL/min at 79% ACN, 0.1% FA, for 7 min, except AP-MS peptides, which were analyzed with a 1h 23 min linear gradient from 2.4% ACN, 0.1% FA to 32% ACN, 0.1% FA, at 0.2 µL/min, 2 min linear gradient to 79% ACN, 0.1% FA, ramping flowrate from 0.3ul/min to 0.4 µL/min, then washed at 79% ACN, 0.1% FA, for 5 min ramping from 0.4 to 0.5 µL/min. For label free phosphoproteomics, SILAC global proteomics, and AP-MS, MS1 scan range is from 350 to 1500 m/z, at resolution 70,000, with Top 15 ions (Top 12 for timecourse) selected for MS2 sequencing at resolution 17,500, normalized collision energy (NCE)=27 after each survey scan. For SILAC phosphoproteomics, MS1 scan range is from 300 to 1750 m/z, at resolution 70,000. Top 12 ions are selected for MS2 sequencing at resolution 35,000, NCE=28 after each survey scan. All MS2 isolation windows are 1.7 m/z with 20 s of dynamic exclusion.

**Maxquant analysis**

Initial timecourse unlabeled phosphoproteomics data were processed together on Maxquant v1.5.1.2 with the following settings: Fixed modifications = “Carbamidomethyl (C), Variable modifications = “Oxidation (M),” “Acetyl (Protein N-term),” and “Phospho (STY),” PSM/Protein FDR = 0.01, min. peptide length = 7, matching time window for matching between runs = 2 min, with 20 min alignment time and all other default parameters (2). Phosphopeptides were searched against the human proteome (Uniprot downloaded 2014/12/3, with 89,706 entries). All SILAC samples were processed together on Maxquant v1.6.0.16 with the same settings, except min. peptide length = 6, matching time window alignment time = 15 min, and max. missed cleavages = 9 (since RS domains on splice factors contain many repeating arginines). Proteomics and phosphoproteomics were searched against the human proteome (Uniprot downloaded on 2018/3/2, with 93,786 entries). SILAC quantification for global proteomics at the protein level requires 1 minimum razor or unique peptide and uses all unmodified and “Oxidation (M)” and “Acetyl (Protein N-term)” modified peptides. AP-MS samples were also processed together on Maxquant v1.6.2.1with the same settings except Variable modifications = “Oxidation (M),” “Acetyl (Protein N-term),” matching time window alignment time = 20 min. Proteins were searched against the human proteome (Uniprot downloaded on 2017/11/15, with 71,544 entries).

Proteomic quantifications, except for the timecourse study, were further evaluated in Perseus (v. 1.6.2.2), where potential contaminants, reverse dummy sequences, and proteins identified by site alone for protein level quantification are excluded (3). Gene ontology annotations were included to identify splicing related proteins. Two biological replicates were grouped and entries with less than 2 valid quantifications were filtered from the final analysis.

**Kinase Set enrichment analysis**

Entire phosphoproteomic results for MM.1S treated with 18 nM Cfz and MM.1S treated with 10 μM melphalan (see **Supplementary Table S4**) were submitted to kinase set enrichment analysis accessed through KSEAapp R package (https://github.com/casecpb/KSEAapp/) (4). Gene name and phosphosite, along with fold change and associated T-test p-value statistic were input to generate kinase activity scores, listed in **Supplementary Table S5**. Bar graphs in **Figure 2F** show top ranked kinases with at least 5 substrates (kinases with 4 or less were excluded from graph).

**Stress Response Immunoblot**

Investigating stress response in **Fig. S3**, 1E+6 cells/mL of AMO-1 and MM.1S were treated with DMSO or specified amounts of Cfz or melphalan for 24 hr, harvested, washed with PBS and pelleted by centrifugation at 300 rcf for 5min, and the cell pellets frozen in 5E+6 cell aliquots in LN2 then stored in -80°C. Cells are thawed and lysed in 125-250 µl 1X RIPA buffer (Millipore) with 1X HALT protease/phosphatase inhibitor, and sonicated with 3 bursts of 5 seconds ON, 10 seconds OFF, @ 20% amplitude with a tip sonicator (BRANSONIC). Protein concentration was quantified with either BCA protein assay kit (Pierce, 23225) or 660 nm protein assay reagent (Pierce, 22660) and ~25-40 µg lysate is loaded per lane and kept consistent across all lanes. Lysate is separated by SDS-PAGE and transferred to PVDF membrane (EMD Millipore, IPFL00010). Membrane is stained with Ponceau-S to assess consistent protein load, then blocked with 5% BSA (Millipore-Sigma, 2930-100GM) in TBS-Tween buffer. Primary antibodies for probing stress response initiation are anti-PERK (CST, 5683P; RRID:AB_10841299), anti-BiP (CST, 3183S; RRID:AB_10695864), anti-phospho-eIF2α (CST, 3398P; RRID:AB_2096481), anti-eIF2α (CST, 5324S; RRID:AB_10692650), anti-phospho-4EBP1 (CST, 2855S; RRID:AB_560835), and anti-4EBP1 (CST, 9644P; RRID:AB_2097841). Horseradish Peroxidase conjugated antibodies: anti-beta-actin-HRP (CST, 5125S; RRID:AB_1903890), secondary anti-Rabbit F(ab)-HRP (Southern Biotech, 4052-05) and secondary anti-Mouse-HRP (Southern Biotech, 1031-05) were used where appropriate. Post-HRP imaging, phospho-specific immunoblots are stripped and re-incubated with corresponding total protein antibody sequentially from the same membrane. Antibodies for DNA damage response markers are anti-phospho-CHK1 (CST, 2348P; RRID:AB_331212) and anti-phospho-H2AX (CST, 9718P; RRID:AB_2118009). To assay carfilzomib induced caspase cleavage of spliceosome components, MM.1S cells were probed with anti-Caspase3 (CST, 9664T;

RRID:AB_2070042), anti-U2AF65=U2AF2 (Abcam, ab37483; RRID:AB_883338), SF3A1 (Abcam, ab128898); SF3B1 (CST, 14434S).

**SRSF1 phosphorylation Immunoblot**

For SRSF phosphorylation immunoblot in **Fig. S3F**, specific antibodies for phosphorylated SRSF proteins are not commercially available and phosphorylation was determined by gel migration. Cells were gently lysed in 100 mM Tris pH 8.5, 10 mM TCEP, 100 mM NaCl, 1% NP-40 alternative, 0.03 U/mL aprotinin and 1mM PMSF on ice for 30 min. 20% of MM.1S DMSO sample was set aside and treated with calf intestinal phosphatase (NEB, M0290S) for 1 hr at 37°C in 1X Cutsmart Buffer (NEB, B7204S) to benchmark migration of dephosphorylated species. 1X HALT protease/phosphatase inhibitor was immediately added to all other samples and the remainder of the DMSO sample. The nucleus was separated from lysate by centrifugation and all samples were denatured with 4 M urea and 0.1% SDS. Immunoblot with anti-SRSF1 (SCBT, sc-33652; RRID:AB_628248), anti-SRSF3 (SCBT, sc-398541), and anti-SRSF6 (SCBT, sc-57954; RRID:AB_785899). Nuclear fractions were too viscous and ran with a smear, so were not considered.

**RNA-seq library preparation (detailed methods)**

Total RNA was extracted with RNeasy Mini-prep kit (Qiagen, 74104). Cells were lysed on ice and genomic DNA was homogenized mechanically using an 18-gauge needle and syringe. For single-timepoint experiments isolated total RNA was cleaned, concentrated with RNA clean & concentrator (Zymo, R1015). mRNA isolation begins with 3 μg total RNA resuspended in lysis/binding buffer, denatured at 65°C for 2 min and treated with SUPERase In (Invitrogen, AM2694), then bound to equilibrated poly-dT magnetic beads (NEB, S1550S), incubated for 10 min, then subjected to a series of washes, according to manufacturer’s protocol and eluted with 20 μL RNAse-free water at 80°C for 2-3 min. cDNA library of 200-300 bp fragments was constructed with Illumina platform TruSeq indexed adapters using Hyper Prep RNAseq Illumina kit (Kapa, KK8540), starting with at least 50 ng isolated mRNA. cDNA library between 200-400 bp were isolated by TBE-Urea PAGE (Life Tech, EC68852BOX) stained with SYBR

Gold stain (Life Tech, S-11494), imaged on Bio-Rad Chemidoc gel imager, and extracted from the gel by manual excision of bands. The gel pieces are blended by centrifuging through a needle prick hole at the bottom of an eppendorf into a collection tube at maximum speed for 3 min, then heated at 70°C for 10 min in 500 µl 10 mM Tris, pH 8.0 (with occasional vortexing). cDNA that diffused out of the gel matrix was separated from gel particles by spinning through Spin-X concentrator (Corning). cDNA was precipitated with 145 mM NaCl, 15 µg/mL glycogen (Thermo, R0551), 58% isopropanol, incubated in -20°C overnight and centrifuged at 4°C for 45 min. The solution is removed and the precipitated white pellet is gently washed with 900 µl 80% ethanol, centrifuged at maximum speed for 5 min at 4°C. The ethanol is removed and the pellet is dried at room temperature for 10 min, then reconstituted in 5-10 µl 10mM Tris pH 8.0. RNA and DNA quantified at all steps by Nanodrop (Thermo Scientific). cDNA library size and quality were evaluated on a Bioanalyzer 2100 (Agilent) with High Sensitivity DNA Kit (Agilent, 5067-4626), before being submitted for next generation sequencing on a HiSeq4000 (Illumina) at the UCSF Center for Advanced Technologies laboratory.

**RNA-seq data analysis**

Four libraries from timecourse study were aligned with Bowtie v0.12.8 allowing for up to two mismatches (5). Aligned rRNA and tRNA reads were discarded. Remaining transcripts were aligned to known canonical transcripts of human genome draft GRCh37/hg19. All other libraries were aligned with HISAT (v2.1.0) (6). The mapped reads were converted from sequence alignment map format to binary alignment map format using Samtools (v1.3.1 for timecourse samples and v0.1.19 for all others) (7). Transcriptome assembled and abundance quantified for four libraries from timecourse study by in-house C++ scripts, which assign and count unique reads mapping to canonical hg19 transcripts. Only uniquely mapping reads were used for analysis in **Fig. 1B**. All other binary alignment mapped reads were quantified for gene-level expression with HTSeq (v0.7.2) (8). Differential gene expression analysis for single-timepoint response study was performed in R with DESeq2 and all differential expression lists are deposited in Gene Expression Omnibus (GEO, accession: GSE124510) (9).

**Cell fixation and fluorescence imaging**

5E+5 AMO-1cells expressing SRSF1-WT(-mCherry-NLS-3XFLAG) and SRSF1-SD and SRSF1-SA mutants were harvested from cell culture, centrifuged at 300 rcf for 5 min. Media was aspirated and the cell pellet was washed once with 1X PBS, centrifuged again for 5 min at 300 rcf and aspirated. The cell pellet was then resuspended in ~200 µl 4% formaldehyde (Pierce, 28906) and incubated in the dark at room temperature for 1h 15 min to fix cells. After fixation, cells were centrifuged at 500 rcf for 5 min, and formaldehyde is carefully removed. Cell pellet is resuspended in PBS and washed twice by centrifugation at 500 rcf for 5 min with aspiration of supernatant. After second wash, cells are resuspended at 5E+6 cells/mL in PBS and pipetted onto poly-L-lysine (Electron Microscopy Sciences, 19320-B) coated #1.5 coverglass (Fisher, 12541B). A small drop of ProLong gold antifade with DAPI (Cell Signaling, 8961S) is added to the cells on the coverslip and gently mixed by stirring with a pipet tip. The coverglass is mounted on glass slide (Fisher, S95933) and cured at room temperature, overnight.

Cells are imaged on a Zeiss Observer Z1 microscope using 63X plan-apochromat oil immersion objective (NA = 1.40). Filter sets for excitation (ex) and emission (em) wavelengths are ex = 335-383 nm, em = 420-470 nm for DAPI and ex = 538-562 nm, em = 570-640 nm, for mCherry. Images were processed in ImageJ (v.1.48) and scale bar represents 10 µm.

**E7107 splicing assay**

RNA was extracted from AMO-1 and MM.1S with RNeasy Mini-prep kit and 500 ng total RNA was used to reverse transcribe polyA-tail mRNA to cDNA with Verso cDNA synthesis kit (Thermo, AB1453B). Splicing activity was determined by comparing abundance of the mature, spliced form of FBXW5 and MBD4, or the unspliced pre-mRNA of cells between conditions by qPCR with SYBR Green supermix (Bio-Rad, 1725272), in technical triplicate on a StepOne Real-Time PCR system (Applied Biosystems). ΔCt between splice targets and “house-keeping” gene PPIA normalized sample variation, and ΔΔCt reports the fold change difference between untreated (DMSO) and E7107 treated samples in 3 biological replicates.

**Patient ex vivo flow cytometry data analysis**

Flow cytometry data was analyzed with FloJo v.8.8.6 to determine relative abundance of CD138+ cells, with respect to all cells counted. **Fig. S7B** shows an example dot plot of stained patient samples and geometric enclosure used to outline CD138+ cells in both DMSO (0 nM E7107) and 100 nM E7107 samples. Reported percent viability is normalized to amount in DMSO treated samples for each patient. Relative abundance of CD138- cells were counted by excluding CD138+ cells and then considering the relative abundance of live, non-SyTOX green stained cells with respect to all non-CD138+ cells.
